# Defining the chromatin-associated protein landscapes on *Trypanosoma brucei* repetitive elements using synthetic TALE proteins

**DOI:** 10.1101/2025.04.22.649942

**Authors:** Roberta Carloni, Tadhg Devlin, Pin Tong, Christos Spanos, Tanya Auchynnikava, Juri Rappsilber, Keith R. Matthews, Robin C. Allshire

**Affiliations:** Centre for Cell Biology and Institute of Cell Biology, School of Biological Sciences, University of Edinburgh. Edinburgh EH9 3BF, Scotland, UK; Institute of Immunology and Infection Research, School of Biological Sciences, University of Edinburgh, Edinburgh EH9 3FL, Scotland, UK; Institute of Biotechnology, Technische Universität, Gustav-Meyer-Allee 25, 13355 Berlin, Germany; NIBRT, Foster Avenue, Blackrock, Co. Dublin, A94 X099, Ireland; Proteomics Science Technology Platform, The Francis Crick Institute, 1 Midland Road, London NW1 1AT, UK

## Abstract

Kinetoplastids, such as *Trypansoma brucei*, are eukaryotes that likely separated from the main lineage at an exceptionally early point in evolution. Consequently, many aspects of kinetoplastid biology differ significantly from other eukaryotic model systems, including yeasts, plants, worms, flies and mammals. As in many eukaryotes the *T. brucei* genome contains repetitive elements at various chromosomal locations including centromere- and telomere-associated repeats and interspersed retrotransposon elements. *T. brucei* also contains intermediate-sized and mini-chromosomes that harbor abundant 177 bp repeat arrays, and 70 bp repeat elements implicated in Variable Surface Glycoprotein (VSG) gene switching. In many eukaryotes repetitive elements are assembled in specialised chromatin such as heterochromatin, however, apart from centromere- and telomere-associated repeats, little is known about chromatin-associated proteins that decorate these and other repetitive elements in kinetoplastids. Here we utilize affinity selection of synthetic TALE DNA binding proteins designed to target specific repeat elements to identify enriched proteins by proteomics. Validating the approach, a telomere repeat binding TelR-TALE identifies many proteins previously implicated in telomere function. Furthermore, the 70R-TALE designed to bind 70 bp repeats indicates that proteins involved in DNA repair are enriched on these elements that reside adjacent to VSG genes. Interestingly, the 177 bp repeat binding 177R-TALE enriches for many kinetochore proteins suggesting that intermediate-sized and mini-chromosomes assemble kinetochores related in composition to those located on the main megabase chromosomes. This provides a first insight into the chromatin landscape of repetitive regions of the trypanosome genome with relevance for their mechanisms of chromosome integrity, immune evasion and cell replication.

## Introduction

Repetitive sequences are scattered across the genomes of many eukaryotes, where they define various functional chromosomal elements (Bringaud et al., 2009; Feschotte, 2008; Kazazian, 2004; Slotkin & Martienssen, 2007). For example, telomeres are generally composed of TG-rich repeats, added by the reverse transcriptase activity of telomerase, which uses its associated RNA as a template (Li, 2021; Pfeiffer & Lingner, 2013), whereas centromere regions often contain extensive tandem arrays of non-conserved repetitive sequences (Allshire & Karpen, 2008; Miga & Alexandrov, 2021; Sullivan & Sullivan, 2020; Thakur et al., 2021). In many eukaryotes such arrays frequently provide a substrate for constitutive heterochromatin formation through di-/tri-methylation of lysine 9 on histone H3 on resident nucleosomes. In addition, repetitive centromeric repeat arrays are associated with the assembly of specialized nucleosomes containing the centromere-specific histone H3 variant, generally known as CENP-A or cenH3 (Allshire & Karpen, 2008; Talbert & Henikoff, 2020). CENP-A nucleosomes form the foundation for kinetochore assembly, which mediates accurate chromosome segregation (Allshire & Karpen, 2008). Other repetitive sequences, such as transposable elements or their remnants, can alter - or have been co-opted to regulate - the expression of nearby genes (Bourque et al., 2018; Fueyo et al., 2022). In many eukaryotes, heterochromatin forms clusters in the nucleus that are generally located at the nuclear periphery or adjacent to nucleoli (Bizhanova & Kaufman, 2020; van Steensel & Belmont, 2017).

Kinetoplastids represent a distinct branch of protozoan eukaryotes within the Euglenozoa that diverged from the main eukaryotic lineage early during their evolution (Cavalier-Smith, 2010). As a result, kinetoplastids are distinct from most other eukaryotes in which cellular mechanisms are intensively studied, including yeasts, fungi, plants, nematodes, insects, and mammals. Many kinetoplastids are parasites that cause diseases in humans and economically-important livestock. *Trypanosoma brucei*, for example, is prevalent in sub-Saharan Africa, where it is transmitted by tsetse flies and causes human African trypanosomiasis and Nagana in cattle (Morrison et al., 2023). Other kinetoplastid parasites that cause human diseases in the tropics include *Trypanosoma cruzi* (Chagas disease) and *Leishmania spp.* (leishmaniasis) (Stuart et al., 2008). Despite their divergence from most eukaryotes, the genomes of kinetoplastids contain a variety of repetitive sequences. The diploid genome of the commonly used laboratory *Trypanosoma brucei* Lister 427 strain has recently been re-characterized with advanced genome assembly methods. The genome contains two homologs for each of the 11 large chromosomes, ranging in size from 900-4,600 kb, 5-6 intermediate chromosomes and ∼100 minichromosomes (Cosentino et al., 2021; Müller et al., 2018) (Rabuffo et al., 2024).

All *T. brucei* chromosomes are linear, and each end terminates with arrays of telomeric (TTAGGG)_n_ repeats that are added by telomerase (Sandhu & Li, 2017). *T. brucei* exhibits the generally well-defined process of Variable Surface Glycoprotein (VSG) gene switching, which allows a proportion of parasites to evade the host immune system at any given time (Barcons-Simon et al., 2023). Most of the 2,634 detected VSG genes are not expressed and reside in arrays in sub-telomeric regions, with others residing on minichromosomes. Only one VSG gene is expressed at any time, and only from one of the estimated fifteen telomere adjacent bloodstream expression sites (BES) (Cosentino et al., 2021). The non-expressed VSG genes provide a library of potential alternative VSGs, so that the parasite has almost limitless potential to vary its protective coat. Further variation in the expressed VSG protein repertoire can be generated by recombination events between VSG genes and VSG pseudogenes, which comprise approximately 80% of the overall gene repertoire (Mugnier et al., 2015) (Cosentino et al., 2021). Non-expressed VSG genes are exchanged with VSG genes residing in expression sites using recombination-directed processes that act on or near 70 bp repeats residing upstream of the resident VSG gene at each BES (Boothroyd et al., 2009; Thivolle et al., 2021). Apart from telomeric (TTAGGG)_n_ repeats at their ends and one or two VSG genes, mini-chromosomes are comprised of tandem arrays of 177 bp repeats, which are also present on the poorly characterized intermediate-sized chromosomes (Ersfeld, 2011; Sloof et al., 1983)(Figure 1A). The function of these 177 bp repeats is unknown, but mini-chromosomes have been shown to be maintained with high stability through mitotic cell divisions, suggesting that a mechanism is in place to ensure their segregation with fidelity to daughter cells (Ersfeld & Gull, 1997; Wickstead et al., 2003). The main 11 megabase-sized chromosomes have been shown to assemble evolutionarily unconventional kinetochores composed of 25 kinetoplastid kinetochore proteins (KKT1-25) that mediate their accurate mitotic segregation and are distinctly different from those of other eukaryotes (Akiyoshi & Gull, 2014; D’Archivio & Wickstead, 2017; Nerusheva et al., 2019) (Akiyoshi & Gull, 2014). ChIP-seq has shown that, on the main megabase-sized chromosomes, kinetochores assemble on different DNA sequences; on some chromosomes kinetochores coincide with tandem arrays of CIR147 repeats or related repeat elements (Akiyoshi & Gull, 2014; Echeverry et al., 2012; Obado et al., 2007). CIR147 repeats produce non-coding transcripts that are processed by Dicer into siRNAs and loaded into Argonaute/TbAGO1 (Tschudi et al., 2012). In addition, SLAC, and *ingi*-related retrotransposons are dispersed across the *T. brucei* genome (Bringaud et al., 2009) and are also transcribed and processed into Ago1-associated siRNA (Tschudi et al., 2012).

**Figure 1.**
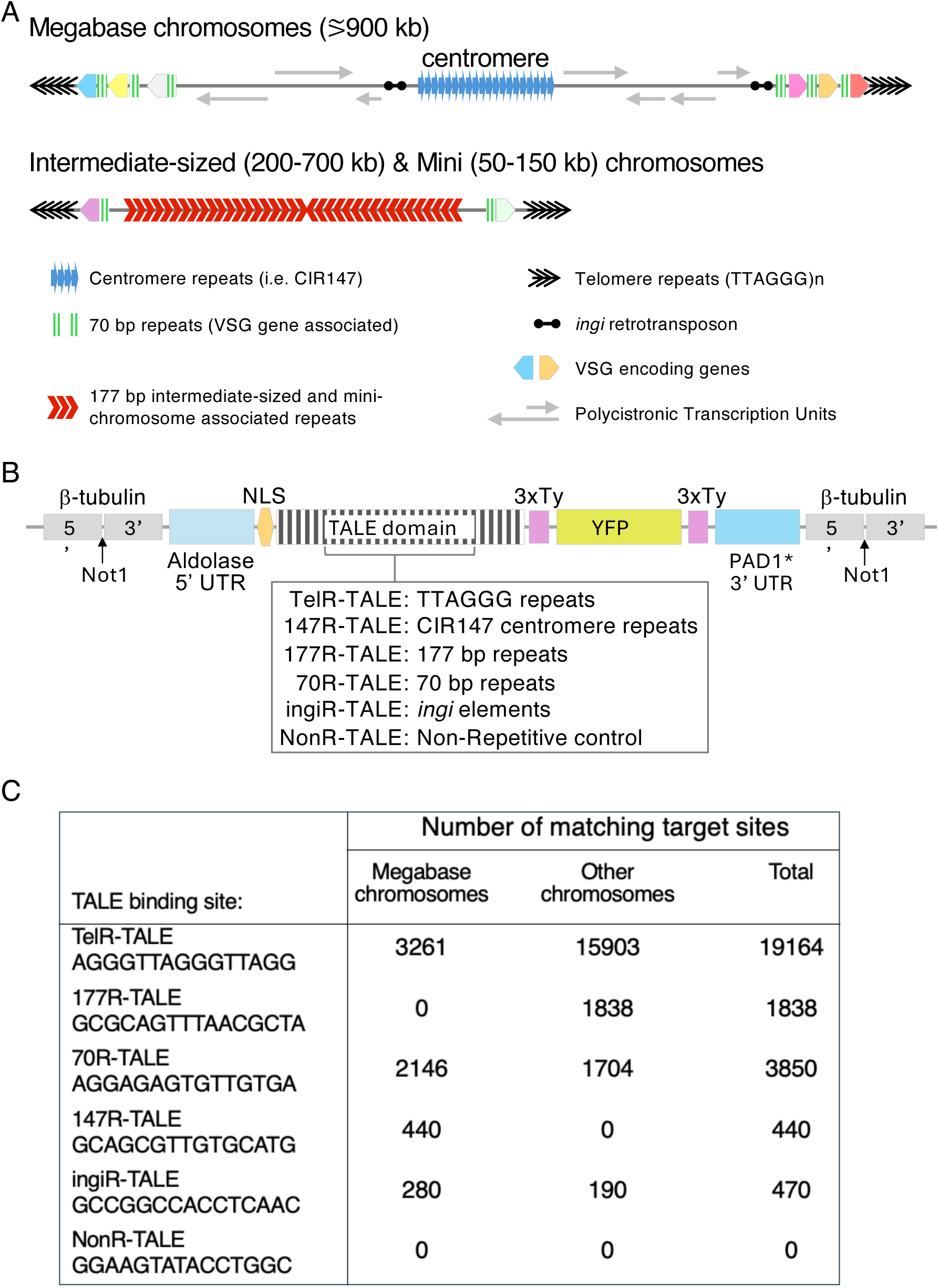
*T. brucei* repetitive elements, TALE design and target site number. **A,** Distinct repetitive elements are present at various locations on *T. brucei* chromosomes. **B.** Construct designed to express the indicated TALE proteins that bind 15 bp target sequences fused to 3xTy1 and YFP tags when integrated at the β-tubulin locus. The Aldolase 5’UTR and PAD1 3’UTR regulate expression levels. A Bleomycin resistance marker gene provides Phleomycin selection (not shown). **C.** Predicted number of target sequences for each TALE in the Lister 427 genome.

The key hallmarks of eukaryotic heterochromatin, di/tri-methylation of histone H3 on lysine 9 (H3K9) or lysine 27 (H3K27), can not be detected in *T. brucei* or other kinetoplastids because their histones, including H3, are particularly divergent rendering useless most existing antibody reagents used for histone post-translational modification (PTM) analyses in other eukaryotes (Deák et al., 2023; Figueiredo et al., 2009). Thus, it is not known which, if any, other modified or unmodified residues on *T. brucei* histones might nucleate repressive chromatin that could be regarded as heterochromatin. However, mass spectrometry has identified a plethora of residues in *T. brucei* and *T. cruzi* histones that exhibit various PTMs (de Lima et al., 2020; Kraus et al., 2020; Maree et al., 2022; Picchi et al., 2017). Some of these PTMs may be involved in forming distinct chromatin structures on repetitive elements through the recruitment of specific proteins analogous to chromodomain protein recruitment via H3K9 or H3K27 methylation in other eukaryotes (Allshire & Madhani, 2018).

To characterise the chromatin context and the possible function of *T. brucei* repetitive elements, we applied an unbiased proteomics-based approach. We exploited synthetic DNA-binding TALE (transcription activator-like effectors) fusion protein expression in *T. brucei* to bind to particular repetitive sequences and, following affinity selection, identify specific factors enriched on these chromosomal regions. Thus, synthetic TALE proteins were designed that were expected to bind the terminal telomeric (TTAGGG)_n_ repeat arrays (TelR-TALE), the most frequent canonical CIR147 centromeric repeat (147R-TALE), core 177 bp repeats (177R-TALE), 70 bp BES-associated repeats (70R-TALE), *ingi-*related retrotransposon repeats (ingiR-TALE) and a Non-Recognised control (NonR-TALE) (Figure 1B). These synthetic TALE proteins were expressed as YFP fusion proteins with a nuclear localisation signal in *T. brucei* Lister 427 Bloodstream form cells with ChIP-seq confirming that they target the repeat elements that they were designed to bind. Validating the approach, affinity purification of TelR-TALE followed by proteomics analyses identified many proteins that were also enriched by affinity purification of the endogenous YFP-tagged *T. brucei* TRF telomere repeat binding protein. Further, several proteins involved in DNA repair-recombination were enriched with affinity purified 70R-TALE suggesting candidates that may be involved in mediating VSG gene switching events via these repeats. Surprisingly, many kinetochore proteins were detected as being enriched on 177 bp repeats. Thus, intermediate-sized and mini-chromosomes may assemble kinetochores and utilise machinery related to that operating on the main eleven chromosomes for their accurate mitotic segregation.

## Results

### Synthetic TALE-YFP fusion proteins that target *T. brucei* repetitive sequences

Five synthetic transcription activator-like effectors TALE proteins were designed that were predicted to specifically bind 15 bp target sequences residing in different repetitive elements using pre-assembled tetramer and trimer modules (Moore et al., 2014) (Figure 1B, C; Fig. S1). BLAST searches confirmed that each selected 15 bp target sequence was unique to the specific target repetitive element with no exact match elsewhere in the *T. brucei* 427 reference genome (Cosentino et al., 2021; Rabuffo et al., 2024). The five TALEs assembled were thus predicted to bind: (i) telomeric (TTAGGG)_n_ repeats residing at all chromosome ends (TelR-TALE) (Blackburn & Challoner, 1984; Van der Ploeg et al., 1984), (ii) the 70 bp repeat arrays that reside upstream of bloodstream VSG gene expression sites, and in shorter tracts adjacent to silent subtelomeric VSG genes and contribute to VSG gene switching events (70R-TALE) (Boothroyd et al., 2009; Glover et al., 2013; Hovel-Miner et al., 2016; Kim & Cross, 2010; Thivolle et al., 2021) (iii) the satellite-like centromere-associated 147 bp Chromosome Internal Repeats (147R-TALE) (Akiyoshi & Gull, 2014; Obado et al., 2005; Tschudi et al., 2012) (iv) the 177 bp satellite repeats that are concentrated on mini- and intermediate-sized chromosomes (177R-TALE) (Wickstead et al., 2004), and (v) a sequence common to the *ingi* clade of non-LTR retrotransposon interspersed repeat elements (ingiR-TALE) (Bringaud et al., 2008). A control NonR-TALE protein was also designed which was predicted to have no target sequence in the *T. brucei* genome. Each synthetic TALE DNA binding domain open reading frame (ORF) was fused at its N-terminus to DNA encoding the *T. brucei* La protein nuclear localization signal (NLS) and at its C-terminus with DNA encoding a 3xTy-YFP tag (Dean et al., 2015; Marchetti et al., 2000). *T. brucei* genes are polycistronic with their expression regulated by RNA processing and turnover. Consequently, the attenuated D1-354 PAD1 3’UTR from the PAD1 gene was placed downstream of each NLS-TALE-3xTy-YFP ORF. Use of this 3’UTR, which drives high level expression in the stumpy transmission stage of parasites but only low level expression in proliferative bloodstream forms (MacGregor & Matthews, 2012) restricted TALE protein expression levels. All constructs carried the Aldolase (ALD) 5’ UTR (ALD) to enable 5’ end RNA processing. Each of the final ALD5’UTR-NLS-TALE-3xTy-YFP*PAD1-3’UTR plasmids was integrated by homologous recombination at the β-tubulin gene locus in monomorphic Lister 427 bloodstream form *T. brucei* cells (for brevity hereon the constructs and proteins produced are refered to as ---R-TALEs; Figure 1B, C; Fig. S1A-C).

Proteins extracted from resultant TelR-TALE, 70R-TALE, 147R-TALE, 177R-TALE, ingiR-TALE, and NonR-TALE *T. brucei* transformants were analysed by anti-GFP and anti-Ty westerns (Figure S2). Cell lines expressing representative TALE-YFP proteins displayed no fitness deficit (Fig. S3A). Five of the six synthetic ORFs produced proteins of the expected size of ∼110 kDa. However, the expression level of NonR-TALE-YFP was lower than other TALE-YFP proteins; this may relate to the lack of DNA binding sites for NonR-TALE-YFP in the nucleus. Moreover, the TelR-TALE protein was smaller than expected; further investigation revealed that the repetitive nature of the telomeric target sequence AGGGTTAGGGTTAGG gave rise to a 612 bp direct repeat within the TALE encoding modules which, following transformation of *T. brucei*, resulted in a deletion event that reduced the predicted recognized target sequence to 8 rather than 15 bases of telomeric repeat (Fig. S1D, E). Nevertheless, about 19,000 copies of the (TTAGGG)_n_ sequence reside at *T. brucei* telomeres and contain the predicted, albeit truncated, TelR-TALE target sequence AGGGTTAG. Indeed further analysis confirmed that the TelR-TALE-YFP protein binds telomeres *in vivo* (see below).

### Synthetic repeat targeting TALE proteins localize to nuclei and are enriched on their cognate sequences

To determine the localisation of the six TALE proteins, anti-GFP immunolocalization was performed on *T. brucei* cells expressing each individual TALE-YFP fusion protein or, as controls, the YFP-TRF telomere (TTAGGG)_n_ binding protein or YFP-KKT2 centromere-associated kinetochore protein (Figure 2; Fig. S3B, Fig S4). All synthetic TALE-YFP proteins and the endogenously tagged YFP-TRF and YFP-KKT2 proteins localized within nuclei, with YFP-TRF and YFP-KKT2 exhibiting distinct nuclear foci as expected for telomeres and centromeres (Li et al., 2005, Akiyoshi & Gull, 2014). The TelR-TALE-YFP and 147R-TALE-YFP localisation patterns were also punctate and comparable to that of YFP-TRF and YFP-KTT2, respectively. Furthermore, the localisation pattern for 177R-TALE-YFP was consistent with the known location of minichromosome 177bp repeats around the nuclear periphery (Ersfeld & Gull, 1997). Both the 70R-TALE-YFP and ingiR-TALE-YFP proteins exhibited a diffuse nuclear signal with no specific sub-nuclear pattern. NonR-TALE-YFP displayed a diffuse nuclear and cytoplasmic signal; unexpectedly the cytoplasmic signal appeared to be in the vicinity the kDNA of the kinetoplast (mitochrondria). We note that artefactual localisation of some proteins fused to an eGFP tag has previously been observed in *T. brucei* (Pyrih et al, 2023).

**Figure 2.**
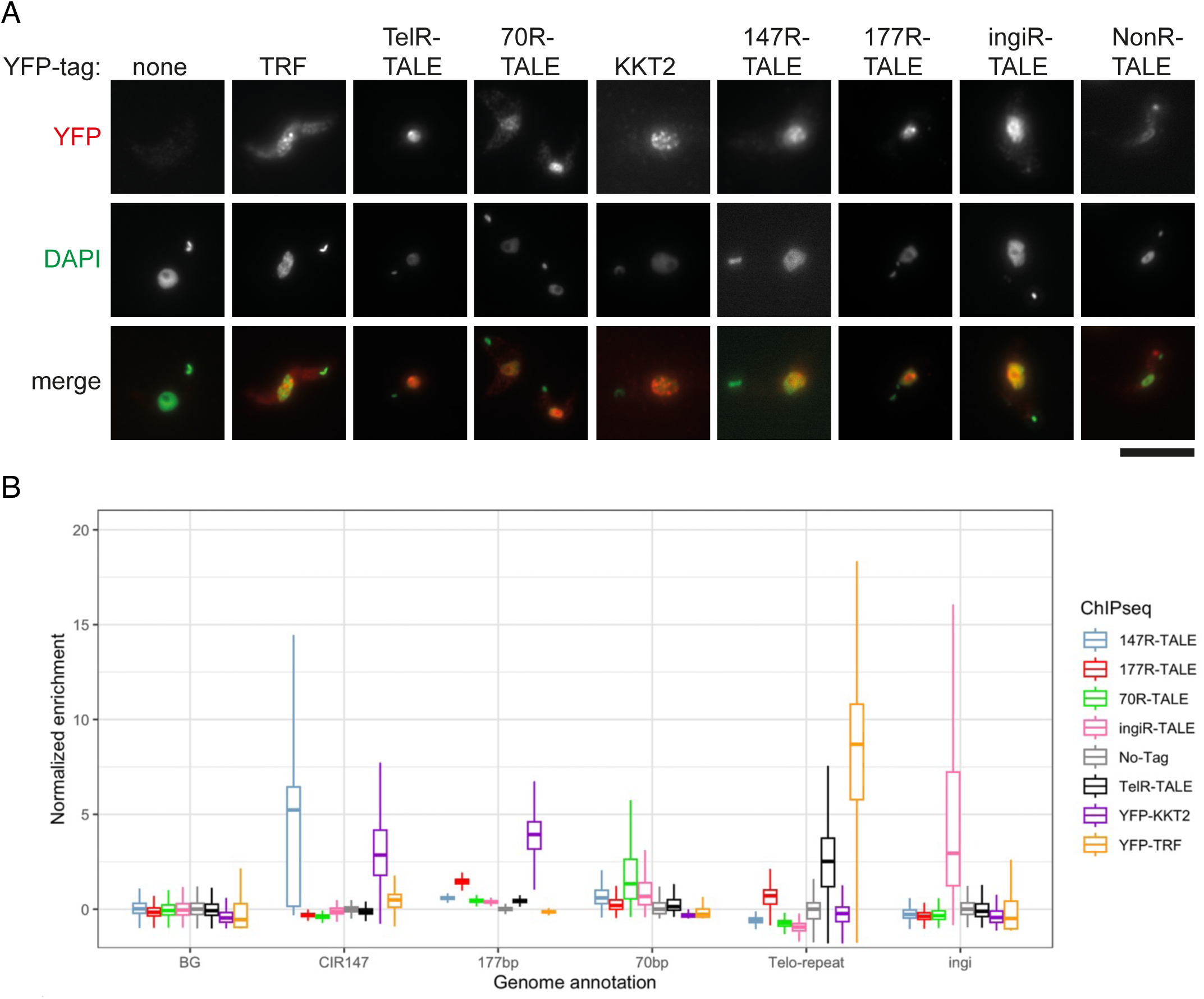
Localization and specific target sequence association of five synthetic TALE-YFP fusion proteins expressed in *T. brucei* compared to YFP-TRF and YFP-KKT. **A.** Bloodstream form Lister 427 *T. brucei* cells expressing the indicated TALE-YFP fusion proteins fixed and TALE-YFP protein localization detected with anti-GFP primary antibody and Alexa Fluor 568-labeled secondary antibody (red). Nuclear and kinetoplast (mitochondrial) DNA were stained with DAPI (green). Control cell expressing telomeric YFP-TRF, centromeric YFP-KKT2 kinetochore protein or wild type Lister 427 cells expressing no YFP, are also shown. Scale bar, 10 μm. **B.** Anti-GFP ChIP-seq analysis for 147R-TALE, 177R-TALE, 70R-TALE, TelR-TALE and ingiR-TALE, demonstrates that each protein is enriched on the repeat elements they were designed to recognize: CIR147 repeats, 177bp repeats, 70bp repeats, telomeric (TTAGGG)n repeats and ingi retrotransposons. Enrichments obtained for the YFP-KKT2 kinetochore protein, the TRF telomere repeat binding protein and with a No-Tag control are shown for comparison. Data are from two biological replicates.

To determine if the TALE proteins were enriched on the repetitive elements that they were designed to bind, anti-GFP ChIP-seq was performed. The resulting ChIP-seq reads were aligned to the most recent *T. brucei* 427 genome assembly (Cosentino et al., 2021; Rabuffo et al., 2024) and the relative specificity compared (Figure 2B). The truncated TelR-TALE protein predicted to bind AGGGTTAG within telomeric (TTAGGG)_n_ arrays was found to be enriched at the end of all megabase-sized, intermediate-sized and mini-chromosomes coincident with telomere repeat binding protein YFP-TRF enrichment (examples, Figure 3B). The 70R-TALE bound to 70 bp repeats that reside upstream of many VSG gene bloodstream expression site loci, regardless of their expression status, 2-8 kb from terminal (TTAGGG)_n_ telomere repeat arrays (Hertz-Fowler et al., 2008) (binding at active BES1 and inactive BES5 is shown in Figure 3B). The ingiR-TALE protein was enriched over the 470 matching *ingi* element target sites dispersed across the *T. brucei* genome and, as expected, these included the region of similarity in RIME, SIDER and DIRE retrotransposons (Bringaud et al., 2008) (Fig. S5).

**Figure 3.**
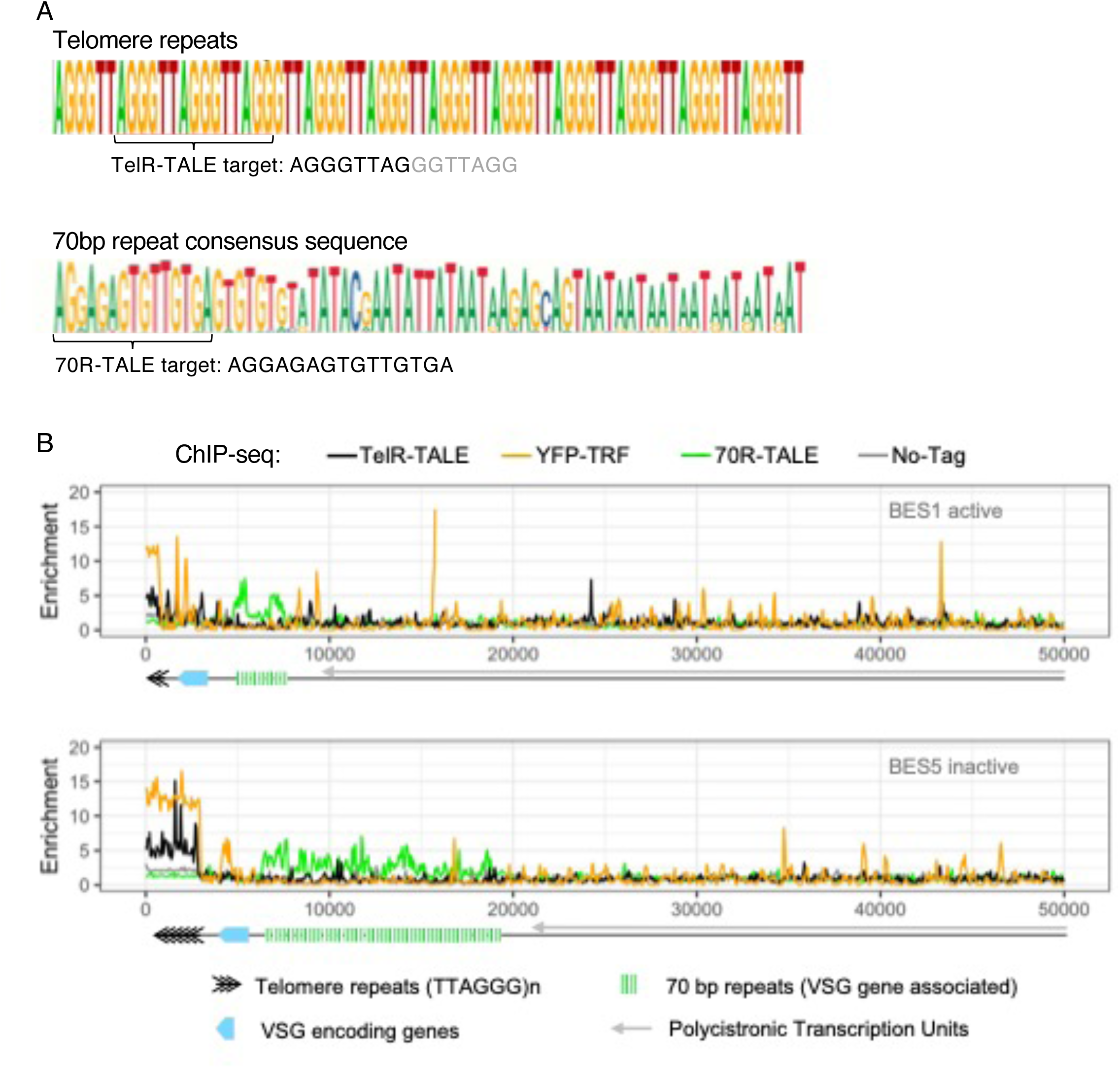
TelR-TALE-YFP and 70R-TALE-YFP are enriched at or near telomeric *T. brucei* blood stream expression sites. **A.** Telomeric repeat (TTAGGG)_n_ sequence (top) and 70bp repeat consensus sequence (bottom). Sequences that TelR-TALE and 70R-TALE were designed to bind is indicated. Deletion of TelR-TALE recognition modules following integration in *T. bruce*i results in recognition of AGGGTTAG rather than the full 15 bp target sequence. **B.** Anti-GFP ChIP-seq for cells expressing TelR-TALE-YFP, YFP-TRF or 70R-TALE–YFP proteins, or 427 cells expressing no YFP tagged protein. Anti-GFP ChIP-seq enrichment profiles are shown for telomeric blood stream expression sites (BES) BES1 (top) and BES5 (bottom). Diagrams show the position of telomeric (TTAGGG)_n_ repeats (black chevrons), VSG genes (blue) and upstream 70 bp repeats (green bars). Data are from two biological replicates. Y axis: Log_2_ values, X axis: base pairs.:

The centromere region of *T. brucei* chromosomes 4, 5, and 8, contain extensive arrays of canonical CIR147 repeats. Divergent but related repeats are associated with the centromeres of the other main chromosomes but no CIR147-related centromere repeats reside on the intermediate-sized or mini-chromosomes. Hence, 147R-TALE, which was designed to bind the TTGACGTGAAAATAC sequence within the consensus CIR147 repeat (Figure 4A), and for which homologous siRNAs are produced (Patrick et al., 2009; Tschudi et al., 2012;), showed enrichment on the cognate repeat arrays at centromeres 4, and 5, and to some extent centromere 8, that are also occupied by the YFP-KKT2 kinetochore protein (Figure 4B, C). In contrast, the 147R-TALE did not decorate the CIR147-related repeats residing at centromeres 9, 10 and 11 or the more divergent classes of repeats bound by YFP-KKT2 at centromeres 1, 2, 3, 6 and 7 (Figure 4C). ChIP-seq analysis for the 177R-TALE showed that this synthetic protein was enriched on target intermediate-sized and mini-chromosome 177bp repeat arrays (Figure 5).

**Figure 4.**
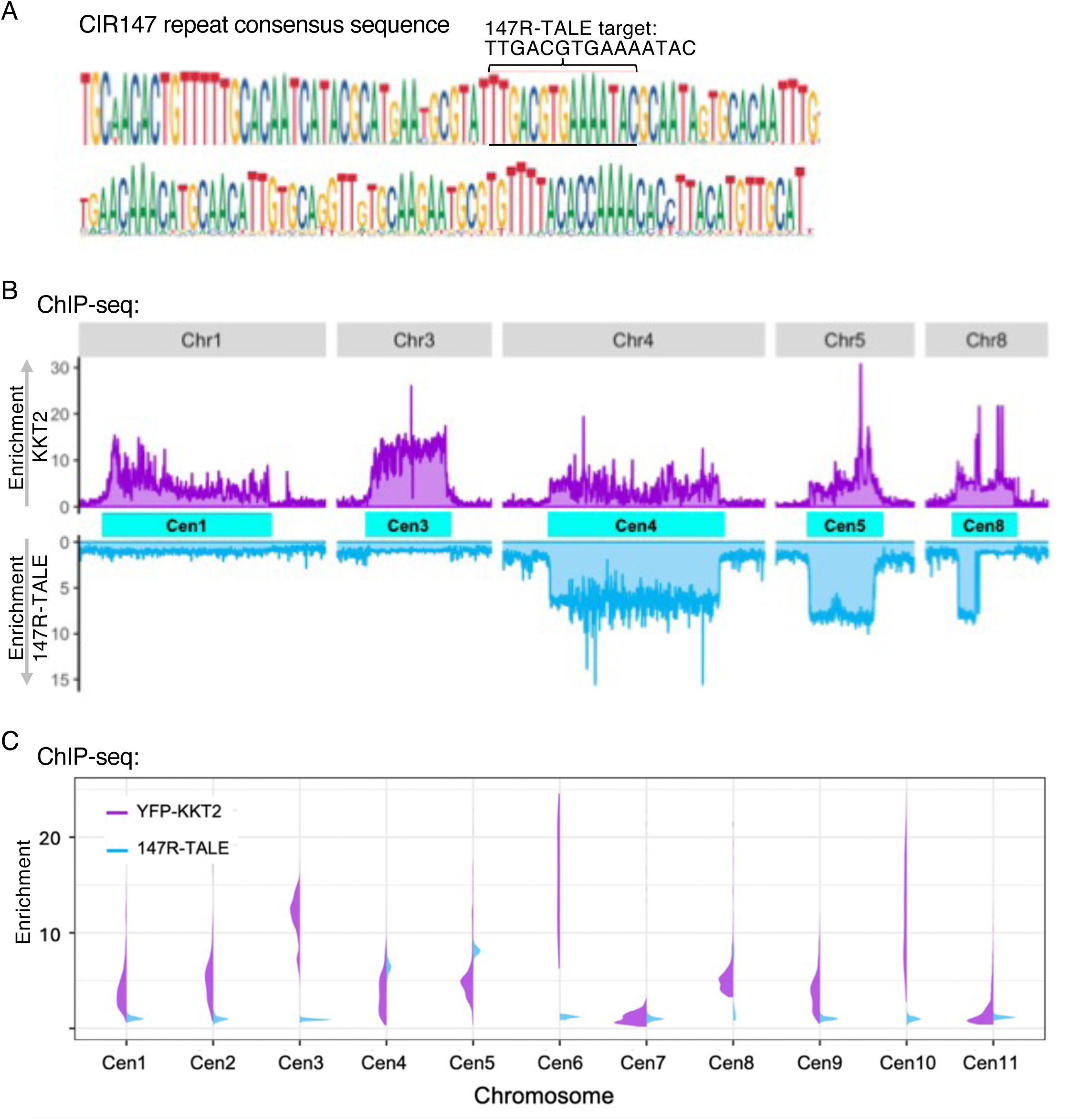
The 147R-TALE-YFP protein is enriched at a subset of centromeres containing canonical CIR147 repeats. **A.** CIR147 repeat consensus sequence. Sequence that 147R-TALE-YFP was designed to bind is indicated. **B.** Comparison of sequences enriched in YFP-KKT2 (purple) and 147R-TALE-YFP (blue) anti-GFP ChIP-seq for chromosomes 1, 3, 4, 5 and 8. DNA from all centromeres are enriched in YFP-KKT2 anti-GFP ChIP-seq whereas only CIR147 repeats at centomeres on chromosomes 4, 5 and 8 are enriched in 147R-TALE-YFP anti-GFP ChIP-seq. **C.** Split-Violin plot demonstrating the relative enrichment of YFP-KKT2 (purple) and 147R-TALE-YFP (blue) over the eleven main chromosome centromere regions. Data are from two biological replicates. Y axis: Log_2_ values.

**Figure 5.**
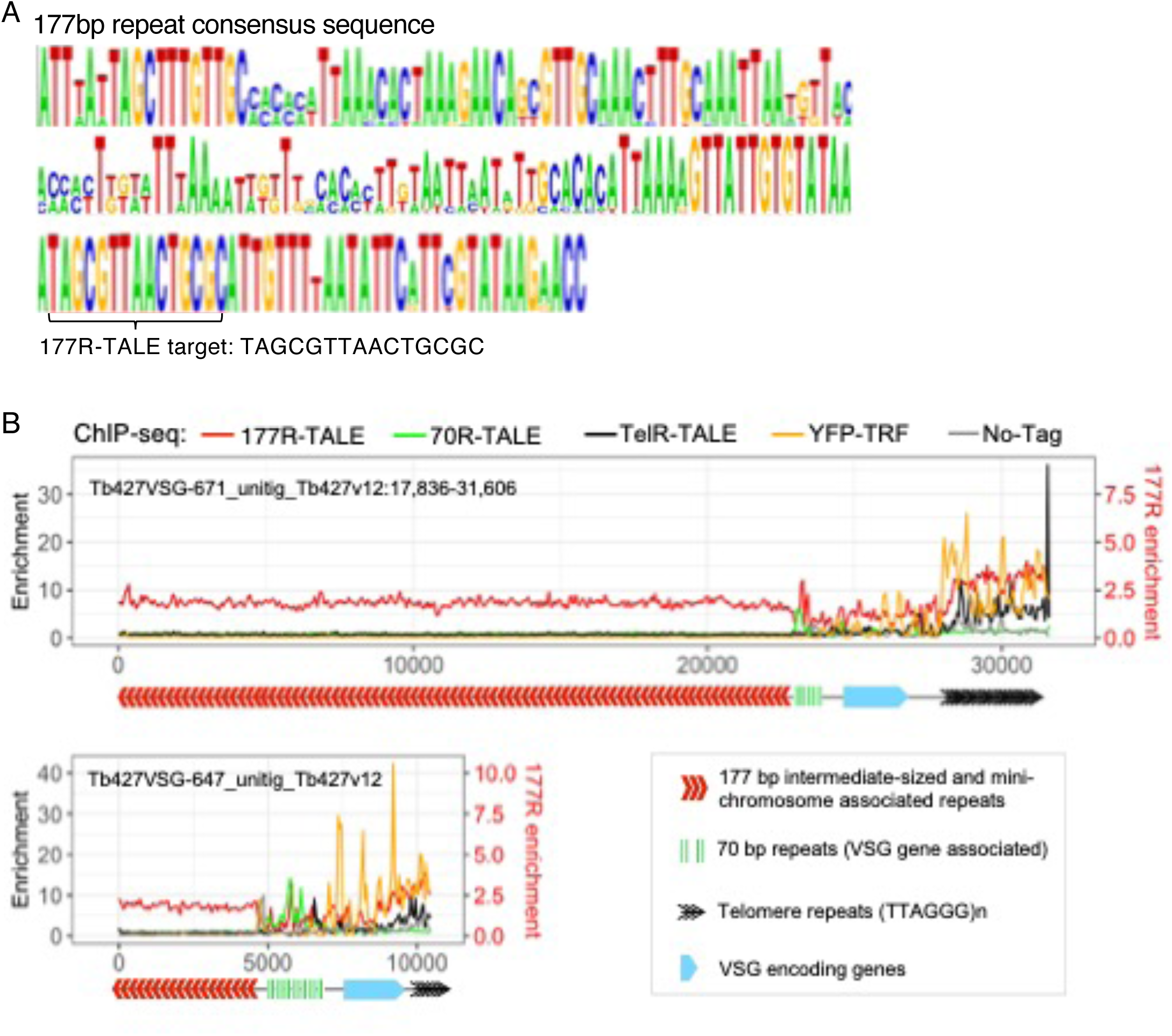
The 177R-TALE-YFP is enriched over 177bp repeats located on intermediate-sized and mini-chromosomes. **A.** 177 repeat consensus sequence. Sequence that 177R-TALE-YFP was designed to bind is indicated. **B.** Distribution of 177R-TALE-YFP, TelR-TALE-YFP, YFP-TRF and 70R-TALE-YFP, at two intermediate/mini-chromosome telomeres determined by anti-GFP ChIP-seq. Anti-GFP ChIP-seq of 427 cells expressing no tagged protein is included as control. Diagrams below ChIP-seq profiles indicate the positions of 177bp repeats (red chevrons), 70bp repeats (green bars), VSG encoding genes (blue) and telomere (TTAGGG)n repeats (black chevrons) within Tb427VSG-671_unitig_Tb427v12:17,836-31,606 (31kb) and Tb427VSG-647_untig_Tb427v12 (10kb). Data are from two biological replicates. Y axis: Log_2_ values, X axis: base pairs.

### TelR-TALE affinity purification verifies use of TALEs to identify repetitive element associated proteins

All five synthetic TALE-YFP proteins were found to target the repetitive elements to which they were designed to bind when expressed in *T. brucei*. At least five proteins have previously been shown to be specifically enriched with the *T. brucei* TRF telomere binding protein in affinity purifications: TIF2, TelAP1, TelAP2, TelAP3 and PolQ/PolIE (Leal et al., 2020; Reis et al., 2018; Weisert et al., 2024). Therefore, to test if repeat-targeted TALE-YFP proteins could be used to identify proteins associated with repetitive elements, we affinity purified solubilised TelR-TALE bound chromatin and compared the associated proteins with those we detected as being enriched with YFP-TRF by mass spectrometry (AP-LC-MS/MS; Figure 6A, B; Fig. S6; Tables S1, S2). As expected known telomere-associated proteins TRF (Tb927.10.12850), TIF2 (Tb927.3.1560), TelAP1 (Tb927.11.9870), TelAP2 (Tb927.6.4330), TelAP3 (Tb927.9.4000), RAP1 (Tb927.11.370) and PolQ/PolIE (Tb927.11.5550) were enriched with affinity purified YFP-TRF (Figure 6A, Fig. S6; Table S1). In addition, replication/repair proteins RPA2 (Replication Factor A; Tb927.11.9130) and PPL2 (PrimPol-Like protein 2; Tb927.10.2520) were also enriched with YFP-TRF, along with the RNA binding proteins ZC3H39 (Tb927.10.14930) and ZC3H40 (Tb927.10.14950), HDAC3 (Tb927.2.2190) and histones (Figure 6A). Using the same affinity selection procedure an overlapping set of 108 proteins were found be enriched with TelR-TALE-YFP bound chromatin (Figure 6B, Fig. S5, Table S2); these included TRF, TIF2, TelAP1, TELAP2, TELAP3, PolQ/PolIE, PPL2, HDAC3 and histones, however, RAP1 was only weakly enriched. In addition, all three Replication Factor A subunits (RPA1, 2, 3; Tb927.11.9130, Tb927.5.1700, Tb927.9.11940) were enriched with TelR-TALE, but only RPA2 with YFP-TRF. Notably, two RNA-associated proteins PABP2 (Tb927.9.10770) and MRB1590 (Tb927.3.1590), which were previously identified as potential telomere-associated proteins (Reis et al., 2018; Weisert et al., 2024), were detected in both our YFP-TRF and TelR-TALE affinity purifications. Moreover, the ZC3H39 (Tb927.10.14930) and ZC3H40 (Tb927.10.14950) RNA binding proteins, that heterodimerize to regulate respiratome transcript levels (Trenaman et al., 2019) were enriched in affinity selections of both proteins Figure S6.

**Figure 6.**
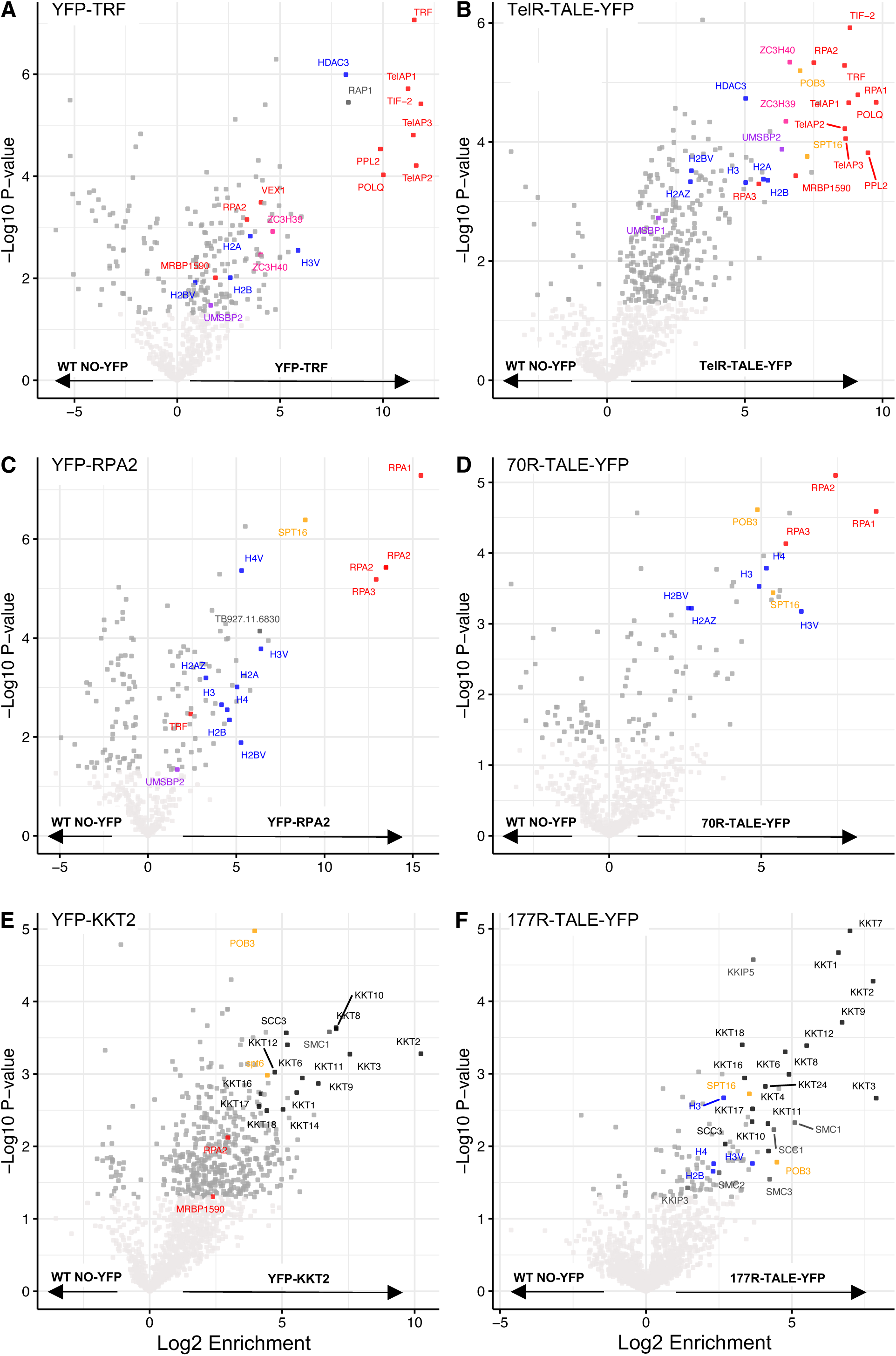
Affinity selection of TelR-TALE-YFP enriches for telomere-associated proteins and 177R-TALE-YFP protein enriches for kinetochore proteins. Affinity selection was performed on control cells expressing YFP-TRF (**A**), YFP-RPA2 (**C**), YFP-KKT2 (**E**) or No-YFP tagged protein and cells expressing synthetic TelR-TALE-YFP (**B**), 70R-TALE-YFP (**D**), 177R-TALE-YFP (**F**). Enriched proteins were identified and quantified by LC-MS/MS analysis relative to the No-YFP tag control. The data for each plot is derived from three biological replicates. Cutoffs used for significance: *P*< 0.05 (Student’s *t*-test). Enrichment scores for proteins identified in each affinity selection are presented in Supplementary Tables (Excel files).

Overall, these data indicate that a core set of known telomere/TRF-associated proteins were also enriched with the synthetic TelR-TALE telomere binding protein. Thus, we conclude that our other synthetic TALE-YFP proteins designed to bind distinct repetitive elements could allow identification of proteins specifically residing on those other sequences *in vivo*. Moreover, a similar set of enriched proteins was identified in TelR-TALE-YFP affinity purifications whether compared with cells expressing no YFP fusion protein (No-YFP), the NonR-TALE-YFP or the ingiR-TALE-YFP as controls (Fig. S7B, S8A; Tables S3, S4, S15). Thus, the most enriched proteins are specific to TelR-TALE-YFP-associated chromatin rather than to the TALE-YFP synthetic protein module or other chromatin.

### Target sequence copy number may determine effectiveness of TALE-YFP proteins in identifying repeat-associated proteins

We estimated that the most recent *T. brucei* 427 genome assembly contains 19164 copies of the telomeric AGGGTTAG target sequence which the truncated Tel-TALE-YFP is predicted to bind within (TTAGGG)_n_ repeat arrays (Cosentino et al., 2021; Rabuffo et al., 2024). In contrast, there are only 440 and 470 targets matching the predicted binding sites TTGACGTGAAAATAC and GCCGGCACCTCAAC for the 147R-TALE and ingiR-TALE synthetic proteins, respectively (Figure 1C). NonR-TALE is predicted to have no matching binding sites in the *T. brucei* TREU 427 genome. To determine if proteins associated with such low copy number TALE-YFP target sequences could be identified we applied the same AP-LC-MS/MS proteomics procedure to *T. brucei* cells expressing 147R-TALE, ingiR-TALE or NonR-TALE. Comparison of either 147R-TALE or ingiR-TALE affinity purifications results with the No-YFP or NonR-TALE-YFP control affinity purifications showed no specific enrichment of any proteins of obvious potential functional interest with either 147R-TALE or ingiR-TALE (Figs. S7E, F; S9; Tables S5-S8). Thus, the nuclear ingiR-TALE-YFP provides an additional chromatin-associated negative control for affinity purifications with the TelR-TALE-YFP, 70R-TALE-YFP and 177R-TALE-YFP proteins (Fig. S8; Tables S15-S17). Moreover, although kinetochore proteins are enriched on CIR147 repeats (Figure 4B, C; Akiyoshi and Gull 2014) no kinetochore proteins were detected in 147R-TALE affinity purifications. Thus, although ChIP-seq showed that both 147R-TALE and ingiR-TALE were enriched on their cognate target sequences it appears that there are insufficient copies of these repeats for our AP-LC-MS/MS procedure to reveal associated proteins above background. We therefore focussed our attention on the 70R-TALE and 177R-TALE synthetic proteins for which there are 3850 and 1828 predicted binding sites in the genome, respectively (Figure 1C).

### The RPA complex is enriched with synthetic 70 bp repeat binding protein

70R-TALE affinity purifications showed enrichment of all three subunits of the Replication Protein A complex (RPA1, RPA2 and RPA3) comparable to the enrichment detected in affinity purification of YFP-RPA2 itself (Figure 6C, D; Table S9, S10). Proteins identified as being enriched with 70R-TALE-YFP (Figure 6D) were similar in comparisons with either the No-YFP, NonR-TALE-YFP or ingiR-TALE-YFP as negative controls (Fig. S6C, S8B; Tables S11, S16). Along with the RPA complex, FACT subunits (SPT16 and POB3), histones and Tb927.11.6830 were also enriched with both YFP-RPA2 and 70R-TALE-YFP affinity purification. This collection of proteins were also enriched in affinity purifications of TelR-TALE, which binds terminal telomeric (TTAGGG)_n_ repeats. In contrast, the 70R-TALE targets 70bp repeats residing several kilobase pairs internal from telomeres (ChIP-seq, Figure 3B). Given the known role for the RPA complex in DNA repair and replication it may have distinct roles in mediating specific DNA transactions via 70 bp repeats and in telomere repeat dynamics (Boothroyd et al., 2009; Li, 2023).

### Kinetochore proteins are enriched on 177 bp repeats bound by 177R-TALE

In contrast to 70R-TALE and TelR-TALE, affinity selection of the 177R-TALE resulted in enrichment of a distinct set of proteins which unexpectedly included 14 of the 25 known kinetoplastid core kinetochore proteins (KKT1, KKT2, KTT3, KKT4, KKT6, KKT7, KKT8, KKT9, KKT10, KKT11, KKT12, KKT16, KKT17, KKT24 (Akiyoshi & Gull, 2014; D’Archivio & Wickstead, 2017; Nerusheva et al., 2019) (Figure 6F; Table S12). The same kinetochore proteins were enriched regardless of whether the 177R-TALE proteomics data was compared with No-YFP, NonR-TALE or ingiR-TALE controls (Fig. S6D, S8C; Tables S13, S17). For comparison YFP-KKT2 was affinity selected from *T. brucei* cells expressing endogenous N-terminal YFP-tagged KKT2 (Figure 6E; Table S14). A clearly overlapping set of proteins was detected in both 177R-TALE-YFP and YFP-KKT2 affinity purifications (Fig. S10). Moreover, the outer kinetochore-associated proteins KKIP3, and KKIP5, which transiently associate with *T. brucei* kinetochores through Aurora B kinase regulation, were also enriched with 177R-TALE, along with Aurora B kinase itself (Table S12) (D’Archivio & Wickstead, 2017; Nerusheva et al., 2019; Zhou et al., 2019). Cohesin complex subunits were also present in both 177R-TALE-YFP and YFP-KKT2 affinity purifications underscoring their expected role in mediating sister-kinetochore cohesion during mitosis. Thus, many KKT kinetochore proteins were found to be enriched with affinity purified 177R-TALE which ChIP-seq showed associates with 177bp repeats but not KKT2-bound centromere regions on the eleven megabase-sized main chromosomes (i.e. 177R-TALE is not enriched on CIR147 or other centromeric repeats; Figure 2B). This finding suggests that kinetochores with a related composition assemble on all *T. brucei* chromosomes regardless of their size classification and that 177 bp repeats attract kinetochore proteins to the intermediate-sized and mini-chromosomes. To explore this possibility further our YFP-KKT2 and 177R-TALE ChIP-seq data was compared over two regions from intermediate-sized chromosomes spanning 177bp repeats to the telomere (Figure 7A). The resulting analysis revealed that both YFP-KKT2 and 177R-TALE proteins are enriched on 177 bp repeat arrays but not adjacent non-repetitive sequences. In contrast, YFP-KKT2 and 147R-TALE, but not 177R-TALE, were enriched over main chromosome centromeric 147 bp repeat arrays (Figure 7B). Taking into account the relative number of CIR147 and 177 bp repeats in the current *T. brucei* genome (Cosentino et al., 2021; Rabuffo et al., 2024), comparative analyses demonstrated that YFP-KKT2 is enriched on both CIR147 and 177 bp repeats (Figure 7C). We conclude that kinetochore proteins assemble on at least a proportion of the individual units within the 177 bp repeat arrays on intermediate-size and/or mini-chromosomes.

**Figure 7.**
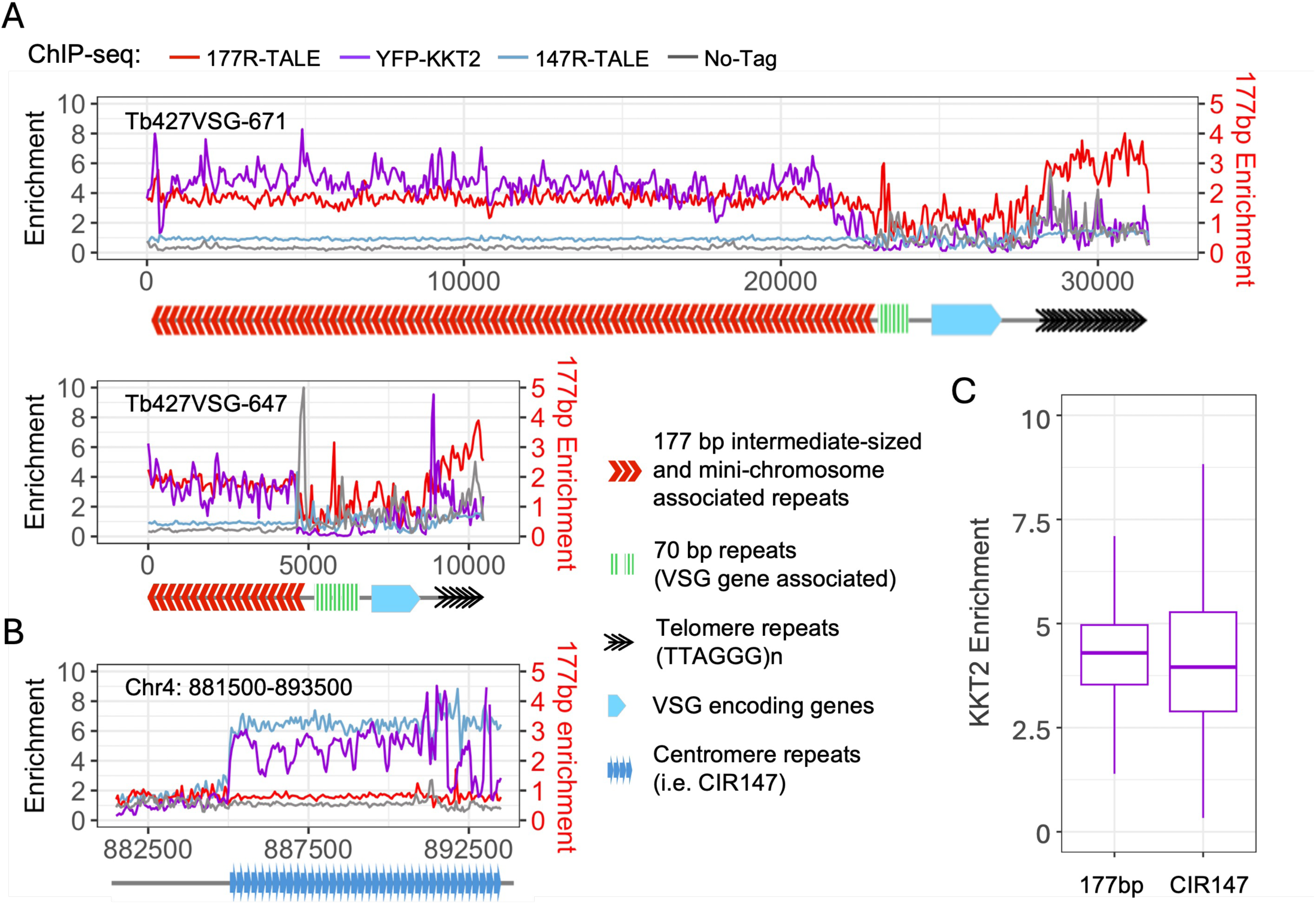
Synthetic 177R-TALE-YFP and YFP-KKT2 kinetochore proteins co-localize over 177bp repeats located on intermediate-sized and mini-chromosomes but not over centromeic CIR147 repeats where 147R-TALE-YFP binds. **A.** Distribution of 177R-TALE, YFP-KKT2 and 147R-TALE over two intermediate/mini-chromosome telomeres determined by anti-GFP ChIP-seq. Anti-GFP ChIP-seq of *T. brucei* 427 cells expressing no tagged protein is included as control. Diagram below ChIP-seq profiles indicates the positions of 177bp repeats (red chevrons), 70bp repeats (green bars), VSG encoding genes (blue) and telomere (TTAGGG)_n_ repeats (black chevrons) within Tb427VSG-671_unitig_Tb427v12:17,836-31,606 (31kb) and Tb427VSG-647_untig_Tb427v12 (10kb). **B.** Comparison of distribution of 177R-TALE, 147R-TALE and YFP-KKT2 over the chromosome 4 CIR147 centromere repeat array and adjacent unique sequences. Chr4:880,000-895,000 (15Kb) and Tb427VSG-671_unitig_Tb427v12:12,000-27,000 (31kb). Diagram below ChIP-seq indicates position of CIR147 repeats. **C.** Comparison of YFP-KKT2 kinetochore protein enrichment on 177 bp and 147 bp repeats. Data are from two biological replicates. Y axis: Log_2_ values, X axis: repeat type.

## Discussion

Repetitive elements are a feature of most eukaryotic genomes with major roles in defining centromeres and telomeres, and influencing the expression of nearby genes through the formation of specific types of chromatin (Allshire & Karpen, 2008; Allshire & Madhani, 2018). Kinetoplastids represent a very distinct early-branching eukaryotic lineage which has highly divergent histones (Alsford & Horn, 2004; Deák et al., 2023; Saha et al., 2021). Consequently little is known about the repertoire of proteins that associate with chromatin formed on repetitive elements in these organisims despite their importance in chromosome segregation, telomere maintenance, and immune evasion. Here we have developed an approach which utilizes a collection of synthetic DNA binding proteins designed to bind telomeric (TTAGGG)_n_ (TelR-TALE), centromeric CIR147 (147R-TALE), ingi-related (ingiR-TALE) dispersed, VSG gene associated 70 bp (70R-TALE) and mini/intermediate chromosome specific (177R-TALE) repeats when expressed in bloodstream form *T. brucei* cells. ChIP-seq demonstrated that all five TALE-based synthetic proteins target the repetitive elements that they were designed to bind. Affinity selection of the TelR-TALE, 70R-TALE and 177R-TALE proteins identified specific sets of enriched proteins. However, the 147R-TALE and ingiR-TALE failed to identify any enrichment of specific proteins following their affinity selection. Encouragingly, the proteins identified as enriched with TelR-TALE showed significant overlap with those identified following affinity selection of the (TTAGGG)_n_ telomere binding protein TRF (Fig. S5). All subunits of the RPA complex were highly enriched with the 70R-TALE and many kinetochore proteins were identified in 177R-TALE affinity purifications.

It was initially surprising that no specific proteins were detected as being enriched following either 147R-TALE or ingiR-TALE pulldown. Given that both of these synthetic proteins target their designated target sequence *in vivo* (Figure 2B), it seems likely that this failure is related to the fact that there are fewer target sites for these synthetic proteins to bind to than the TelR-TALE, 70R-TALE and 177R-TALE proteins which clearly identified proteins bound in their immediate vicinity in *T. brucei* cells. Thus, although proteins that bind CIR147 or ingi-related repeats *in vivo* may be present, their level of enrichment may not be sufficient to allow detection above background by proteomic analyses following 147R-TALE or ingiR-TALE pulldown. It is also possible that the binding of the 147R-TALE and ingiR-TALE proteins dislodges a significant proportion of the proteins that normally bind these repeats, thus reducing their enrichment. Regardless, the 147R-TALE and ingiR-TALE proteins were well expressed in *T. brucei* cells, but their affinity selection did not significantly enrich for any relevant proteins. Thus, 147R-TALE and ingiR-TALE provide reassurance for the overall specificity for proteins enriched in TelR-TALE, 70R-TALE and 177R-TALE affinity purifications.

The TelR-TALE binds telomeric (TTAGGG)_n_ repeats *in vivo* and copurifies with a collection of proteins known to function at trypanosome telomeres, therefore demonstrating that a synthetic protein designed to bind a repetitive element can be used to identify other proteins enriched over those repeats (Figure 6A, B and Fig. S5). Apart from known telomere binding proteins, the two zinc finger proteins ZC3H39 (Tb927.10.14930) and ZC3H40 (Tb927.10.14950) were enriched in both TelR-TALE-YFP and YFP-TRF affinity selections. The fact that both ZC3H39 and ZC3H40 were enriched by affinity selection with these independent baits − one endogenous (YFP-TRF) and the other a synthetic telomere repeat binding protein (TelR-TALE-YFG) − suggests that at least a proportion of these proteins are present near, and may have some function at, telomeres. Although both ZC3H39 and ZC3H40 have also been shown to be involved the post-transcriptional regulation of transcripts encoding respiratory chain proteins and located primarily in the cytoplasm, both were identified in a genome-wide RNAi screen for telomeric gene derepression that also selected the VSG expression site regulator VEX1 (Trenaman et al., 2019). Hence, in addition to their role in respiratory complex gene regulation, ZC3H39 and ZC3H40 might play some additional role in the regulation of gene expression near telomeres. It is possible that they act through telomerase recuitment at telomeres, the regulation of telomere repeat-containing RNA (TERRA) transcripts that are produced at VSG active telomeres (Saha et al., 2021) or engagement of some other regulatory complex associated with telomeres.

Analysis of 70bp repeat associated proteins via specific 70R-TALE affinity selection identified RPA1, 2 and 3. This heterotrimeric complex is enriched at single stranded DNA and assocated with DNA damage and double stranded DNA breaks. The accumulation of these proteins at 70bp repeats is consistent with the function of these sequences in the initiation of recombination events involved in surface antigen switching, with trypanosomes being unusual in not activating a DNA damage cell cycle checkpoint thereby allowing continued proliferation whilst promoting antigenic diversity (Glover et al., 2019). Furthermore, the enrichment of FACT complex components with repeat bound 70R-TALE again highlights 70 bp repeats as an expected focus of recombination events and expression site activity. FACT depletion is known to alleviate repression at these silent VSG expression sites by generating a more open chromatin conformation and reciprocally decreases expression from the active VSG expression site (Denninger & Rudenko, 2014).

Kinetoplastid kinetochores are unusual in that they are composed of at least 26 proteins that bear little resemblance to the 40-100 constitutive and transient kinetochore-associated proteins that assemble at conventional eukaryotic centromeres (Akiyoshi & Gull, 2014; Ballmer et al., 2024; D’Archivio & Wickstead, 2017; Yatskevich et al., 2023). In *T. brucei* kinetochores assemble at a single location on both copies of the 11 main diploid chromosomes. The centromeres of chromosomes 4, 5 and 8 contain CIR147 repeat arrays over which kinetochore proteins are enriched while the centromeres of other megabase chromosomes form on less well characterised repetitive elements (Akiyoshi & Gull, 2014; Echeverry et al., 2012; Obado et al., 2007). In addition, the characterisation of the many *T. brucei* mini and intermediate chromosomes remains incomplete due to the presence of long tandem 177 bp repeat arrays. Our ChIP-seq analyses of synthetic 177R-TALE-YFP location showed that it associated with 177 bp repeats *in vivo* but not the adjacent VSG gene regions on mini-chromosomes or any region of the 11 megabase chromosomes (Figure 7). The detection of a plethora of kinetochore proteins on 177R-TALE-YFP bound chromatin indicates that kinetochores or a sub-kinetochore complex also assembles on the 177bp repeats of mini and intermediate *T. brucei* chromosomes. Consistent with this finding some enrichment of YFP-tagged KKT2 and KKT3 was previously detected using a model minichromosome assemblage, and depletion of kinetochore proteins was shown to cause aberrant segregation of mini-/intermediate chromosomes (Akiyoshi & Gull, 2014). If, as previously suggested (Akiyoshi & Gull, 2014), mini- and intermediate-177bp repeat bearing chromosomes segregate by somehow ‘hitching a ride’ via kinotochores that are actually assembled on the main chromosomes then it might be expected that 177R-TALE-YFP ChIP-seq would register some signal over centromere regions of the main chromosomes, however, no such signal was observed (Figures 4, 7; Fig S1). The 17 kinetochore proteins (KKT1, 2, 3, 4, 6, 7, 9, 10, 11, 12, 16, 17, 18, 24, KKIP3, KKIP4) detected in 177R-TALE-YFP affinity purifications represent most of the components considered to comprise the core kinetochore but represent only a subset of the 26 known *T. brucei* main structural kinetochore proteins. It is possible that a more rudimentary kinetochore is assembled on mini and intermediate chromosome 177 repeat arrays and that these are sufficient to mediate their accurate segregation.

Interestingly, although targetting TALE proteins to different repetitive sequences selected components specific to each repeat type, some overlap in the proteins detected was observed. For example, enrichment of telomere-associated proteins was detected in some affinity selected samples using 177R-TALE-YFP, presumably resulting from the juxtaposition of telomeric repeats and 177bp repeats on minichromosomes. Supporting this, KKT3 was reciprocally detected in samples affinity selected using YFP-TRF. Similarly, enrichment of both FACT subunits with 177R-TALE may simply reflect the proximity of silent telomeric chromatin on minichromosomes, or it may also indicate that FACT contributes to a particular chromatin environment at 177 bp repeats.

Although proteins associated with TALE-YFP fusions targetting telomeric, 70bp and 177bp repeats were successfully identified, our analyses suggest that target sequences need to be present in many copies (>1000) in a *T. brucei* genome of ∼35 Mb to successfully identify associated proteins. Thus, the 147R-TALE-YFP which targets 440 copies of the canonical centromeric CIR147 repeat resulted in no enrichment of associated proteins, although other paramaters may also influence the ability of a sequence bound TALE-YFP protein to enrich for nearby chromatin-associated proteins. Such parameters may include the affinity that the target chromatin bound proteins have for the repeat sequence of interest, the relatively low affinity that these chromatin proteins may have for any other chromosomal region and their overall relative abundance (for detailed discussion see Gauchier et al, 2020). Methods such as CUT&RUN (Skene & Henikoff, 2017), which should selectively release only TALE-bound chromatin, followed by affinity selection (similar to CUT&RUN.ChIP (Brahma & Henikoff, 2019) might improve protein enrichment relative to background and allow identification of proteins associated with less abundant sequences. An alternative to synthetic TALE proteins is to utilise tagged catalytically dead Cas9 targetted to specific sequnces via a CRISPR embedded guide RNA. Fusion of TALE or dCAS9 probes to APEX or BirA* enzymes could also be incorporated to perhaps improve the identification of proteins that reside close to the synthetically targeted DNA binding protein (Gao et al., 2018; Myers et al., 2018). Cas9/CRISPR systems allow precise genome editing in *T. brucei* (Rico et al., 2018; Vasquez et al., 2018) such that the development of dCas9-based CRISPR tools may improve the performance of future sequence targeted proteomics. However, an advantage of TALE protein use is that only a single entity needs to be expressed that directy targets the sequence of interest.

In conclusion we have successfully deployed TALE-based affinity selection of proteins associated with repetitive sequences in the trypanosome genome. This has provided new information concerning telomere biology, chromosomal segregation mechanisms and immune evasion strategies employed by these evolutionarily divergent pathogens. As well as providing an orthogonal corroboration of existing knowledge of protein interactions with discrete genomic features, this has provided new entry points to dissect these parasite’s chomatin architecture. We anticipate extension to other kinetoplastid parasites could assist exploration of *Leishmania* genome instability as a response to environmental adaptation where, for example, the highly abundant SIDER family (70 fold more numerous than in *T. brucei*; (Bringaud et al., 2007) might overcome the copy number limitations of analysing retransposon sequences analysed in our study. Likewise, the 195nt satellite DNA in *T. cruzi* represents 5-10% of the parasite genome and is sufficiently abundant to allow analysis of associated proteins (Elias et al., 2003).

## MATERIALS AND METHODS

### TALEs target sequence design

All synthetic TALE proteins were designed to bind 15 bp target sequences following a T/thymine base as required for the TALEN kit (Ding et al., 2013). The design of the minichromosomal 177 bp repeat binding TALE was informed by available sequences (Wickstead et al., 2004). The ingi repeat TALE was designed to bind a target within the conserved 5’ region 79 bp of related transposable elements (Bringaud et al., 2008). For design of the CIR147 binidng TALE published sequences were used as reference (Obado et al., 2007; Patrick et al., 2009), however a CIR147 bp target sequence with only one exact match was picked. TALEs were designed that were predicted predicted to bind the known 70 bp repeats Boothroyd et al., 2009) and terminal (TTAGGG)n repeats. A control NonR-TALE predicted to have no recognised target in the *T. brucei* geneome was designed as follows: BLAST searches were used to identify exact matches in the TREU927 reference genome. Candidate sequences with one or more match were discarded. Each TALE was assembled using the Musunuru/Cowan TALEN kit protocol (Ding et al., 2013) and subsequently placed in a vector that allowed expression in *Trypanosoma brucei* bloodstream cells as described in the main text.

### Trypanosome cell culture

*Trypanosoma brucei brucei* Lister 427 bloodstream form monomorphic cells were used for all experiments. Cell lines were grown at 37°C and 5% CO2 in HMI-9 medium supplemented with 10% Fetal Calf Serum (Gibco), 1% 32 Penicillin-Streptomycin (Gibco) and selective drug(s) when required (Hirumi & Hirumi, 1989). Cell cultures were maintained below 3 x 10^6^ cells/ml. Pleomycin 2.5 μg/ml was used to select transformants containing the TALE construct BleoR gene.

### Trypanosome transfections

5 x 10^7^ cells were harvested per transfection by centrifugation at 1000 g, 10 min. Cells were washed once with 5 ml TbBSF transfection buffer (Schumann Burkard et al., 2011) and pelleted again by centrifugation at 1000 g, 10 min before resuspending in 100 μl ice-cold TbBSF transfection buffer, and transferred to an electroporation cuvette (Ingenio). 10-20 μl of DNA for transfection containing 1-5 μg DNA was added to the cuvette. Cells were electroporated in the Amaxa Nucleofector II (Lonza) using the X-001 programme for bloodstream cells. A “no DNA” mock transfection was always performed in parallel as a negative control. Electroporated bloodstream cells were added to 30 ml HMI-9 medium and two 10-fold serial dilutions were performed in order to isolate clonal Pleomycin resistant populations from the transfection. 1 ml of transfected cells were plated per well on 24-well plates (1 plate per serial dilution) and incubated at 37°C and 5% CO2 for a minimum of 6 h before adding 1 ml media containing 2X concentration Pleomycin (5 μg/ml) per well. A positive control was also performed by adding media containing no selective drug to 12 wells of the control transfection plate.

### Western analyses

Cells were harvested by centrifugation at 1000 g, 10 min, washed with 1X PBS and resuspended in 1X PBS + 4X NuPAGE LDS Sample Buffer (Thermo Fisher Scientific) to give a final concentration of 5×10^6^ cells per 10 μl. Samples were then boiled at 95°C for 5 min to ensure cells were dead before removal from the CAT3 facility. Samples were then subjected to sonication using a Diagenode Bioruptor for 10 cycles, 30s ON/30s OFF at 4°C on high setting to shear the DNA and reduce the viscosity to aid loading on gels. Samples were run on NuPAGE Bis-Tris Mini Gels (Thermo Fisher Scientific) in a Mini Gel Tank (Thermo Fisher Scientific) in 1X NuPAGE MES Running Buffer at 200 V. Following PAGE, proteins were transferred onto nitrocellulose membranes in a Mini Blot Module (Thermo Fisher Scientific) at 20 V for 1 h. Membranes were stained with Ponceau S (Sigma-Aldrich) to assess efficiency of protein transfer. After blocking with 5% milk/PBS-T (PBS + 0.05% tween), membranes were incubated with mouse anti-GFP (Roche) (1:1000 in 5% milk in PBS-T) or anti-BB2 antibody (Hybrydome mouse monoclonal, clone BB2) (1:5 in 5% milk in PBS-T) at 4°C overnight on a lab rocker, then washed with PBS-T and incubated with HRP-conjugated anti-mouse secondary antibody (1:2500 in 5% milk in PBS-T) at room temperature for 1 h. Membranes were washed with PBS-T and incubated with Amersham ECL Prime Western Blotting Detection Reagent (GE Healthcare) following manufacturer’s instructions. Proteins were visualised using Amersham Hyperfilm ECL (GE Healthcare).

### Fluorescent immunolocalization

Cells were fixed with 4% paraformaldehyde for 10 min on ice. Fixation was stopped with 0.1 M glycine. Cells were added to polylysine-coated slides and permeabilized with 0.1% Triton X-100. The slides were blocked with 2% BSA. Rabbit anti-GFP primary antibody (Thermo Fisher Scientific A-11122) was used at 1:500 dilution, and secondary Alexa Fluor-568 or -488 goat antirabbit antibody (Thermo Fisher Scientific) was used at 1:1000 dilution. Images were taken with a Zeiss Axio Imager microscope.

### Chromatin immunoprecipitation and sequencing

3.5 x 10^8^ parasites were fixed with 0.8% formaldehyde for 20 min at room temperature. Cells were lysed and sonicated in the presence of 0.2% SDS for 30 cycles (30 sec on, 30 sec off) using the high setting on a Bioruptor sonicator (Diagenode). Cell debris were pelleted by centrifugation, and SDS in the lysate supernatants was diluted to 0.07%. Input samples were taken before incubating the rest of the cell lysates overnight with 10 μg rabbit anti-GFP antibody (Thermo Fisher Scientific A-11122) and Protein G Dynabeads. The beads were washed, and the DNA eluted from them was treated with RNase and Proteinase K. DNA was then purified using a QIAquick PCR purification kit (Qiagen), and libraries were prepared using NEXTflex barcoded adapters (Bio Scientific). The libraries were sequence on Illumina NextSeq (Western General Hospital, Edinburgh). In all cases, 75-bp paired-end sequencing was performed. Our subsequent analyses were based on two replicates for all TALEs.

### ChIP-seq data analysis

Sequencing data were mapped to the Tb427V12 genome build(Rabuffo et al., 2024) using Bowtie2 (version 2.4.2), with duplicate reads removed using SAMtools (Danecek et al., 2021). The default mode of Bowtie 2 was used, which searches for multiple alignments and reports the best one or, if several alignments are deemed equally good, reports one of those randomly. The peaks were identified using MACS2 (version 2.2.7.1) broad peak call. The ChIP samples were normalized to their respective inputs (ratio of ChIP to input reads) and the genome overview were using deeptools (Ramírez et al., 2016) with 5bp sliding window.

#### Background enrichment calculation

The genome was divided into 50 bp sliding windows, and each window was annotated based on overlapping genomic features, including CIR147, 177 bp repeats, 70 bp repeats, and telomeric (TTAGGG)_n_ repeats. Windows that did not overlap with any of these annotated repeat elements were defined as “background” regions and used to establish the baseline ChIP-seq signal. Enrichment for each window was calculated using bamCompare, as log₂(IP/Input). To adjust for background signal amongst all samples enrichment values for each sample were further normalized against the corresponding No-YFP ChIP-seq dataset.

### Affinity purification and LC-MS/MS proteomic analysis

Cells, 3.5 x 10^8^, were lysed per IP in the presence of 0.2% NP-40 and 150 mM KCl. Lysates were sonicated briefly (three cycles, 12 sec on, 12 sec off) at a high setting in a Bioruptor (Diagenode) sonicator. The soluble and insoluble fractions were separated by centrifugation, and the soluble fraction was incubated for 1 h at 4 C with beads cross-linked to mouse anti-GFP antibody (Roche 11814460001). The resulting immunoprecipitates were washed three times with lysis buffer, and protein was eluted with RapiGest surfactant (Waters) for 15 min at 55 C. Next, filter-aided sample preparation (FASP) (Wiśniewski et al. 2009) was used to digest the protein samples for mass spectrometric analysis. Briefly, proteins were reduced with DTT and then denatured with 8 M urea in Vivakon spin (filter) column 30K cartridges.

Samples were alkylated with 0.05 M IAA and digested with 0.5 μg MS grade Pierce trypsin protease (Thermo Fisher Scientific) overnight, desalted using stage tips (Rappsilber et al. 2007), and resuspended in 0.1% TFA for LC-MS/MS. Peptides were separated using RSLC Ultimate 3000 system (Thermo Fisher Scientific) fitted with an EasySpray column (50 cm; Thermo Fisher Scientific) using 2%–40%–95% nonlinear gradients with solvent A (0.1% formic acid) and solvent B (80% acetonitrile in 0.1% formic acid). The EasySpray column was directly coupled to an Orbitrap Fusion Lumos (Thermo Fisher Scientific) operated in DDA mode. “TopSpeed” mode was used with 3-sec cycles with standard settings to maximize identification rates: MS1 scan range 350–1500 mz, RF lens 30%, AGC target 4.0e5 with intensity threshold 5.0e3, filling time 50 msec and resolution 120,000, monoisotopic precursor selection, and filter for charge states two through five.

HCD (27%) was selected as fragmentation mode. MS2 scans were performed using an ion trap mass analyzer operated in rapid mode with AGC set to 2.0e4 and filling time to 50 msec. The dynamic exclusion was set at 60s.

-

The MaxQuant software platform (Cox and Mann, 2008) version 1.6.1.0 was used to process the raw files and search was conducted against *Trypanosoma brucei brucei* complete/reference proteome (Uniprot - released in April, 2019), using the Andromeda search engine (Cox *et al*, 2011). For the first search, peptide tolerance was set to 20 ppm while for the main search peptide tolerance was set to 4.5 pm. Isotope mass tolerance was 2 ppm and maximum charge to 7. Digestion mode was set to specific with trypsin allowing maximum of two missed cleavages. Carbamidomethylation of cysteine was set as fixed modification. Oxidation of methionine was set as variable modification. Label-free quantitation analysis was performed by employing the MaxLFQ algorithm as described by Cox *et al* (2014). Absolute protein quantification was performed as described in Schwanhäusser *et al*, 2011. Peptide and protein identifications were filtered to 1% FDR. Statistical analysis and visualisation were performed using Perseus ver. 1.6.2.1 (Tyanova, *et al,* 2016).

### NGS and Proteomics Data Access

The sequencing data generated in this study will be accessible on the NCBI Gene Expression Omnibus (GEO; https://www.ncbi.nlm.nih.gov/geo/) via accession number GSE295698.

The proteomics data generated in this study can be accessed on the Proteomics Identification Database (PRIDE; https://www.ebi.ac.uk/pride/) with accession number PXD063130

## Acknowledgements

The authors thank Alison Pidoux for comments on the manuscript and assistance with images, Bungo Akiyoshi for comments and discussion and Shaun Webb of the Wellcome Centre for Cell Biology and Discovery Research Platform for Hidden Cell Biology Bioinformatics Core for maintaining servers and pipelines for processing sequencing data.

The authors also thank Julie Young for laboratory management support for trypanosome culture during this project. This work was funded by a UKRI/BBSRC EastBio PhD studentship supporting Tadhg Devlin (BB/M010996/1), an MRC Research Grant awarded to R.C.A and K.R.M and supporting R.C. (MR/T04702X/1), a Wellcome Investigator Award to K.R.M. (221717), a Wellcome Principal Research Fellowship to R.C.A. supporting R.C. and T.A (200885; 224358), a Wellcome Instrument grant to J.R. (108504), and core funding for the Wellcome Centre for Cell Biology (203149) and subsequently the Wellcome funded Discovery Research Platform for Hidden Cell Biology DRP-HCB supporting C.S. (226791).

For the purpose of Open Access, the authors have applied a CC BY public copyright licence to any Author Accepted Manuscript version arising from this submission.

## Supplementary Figure Legends

**Figure S1.**
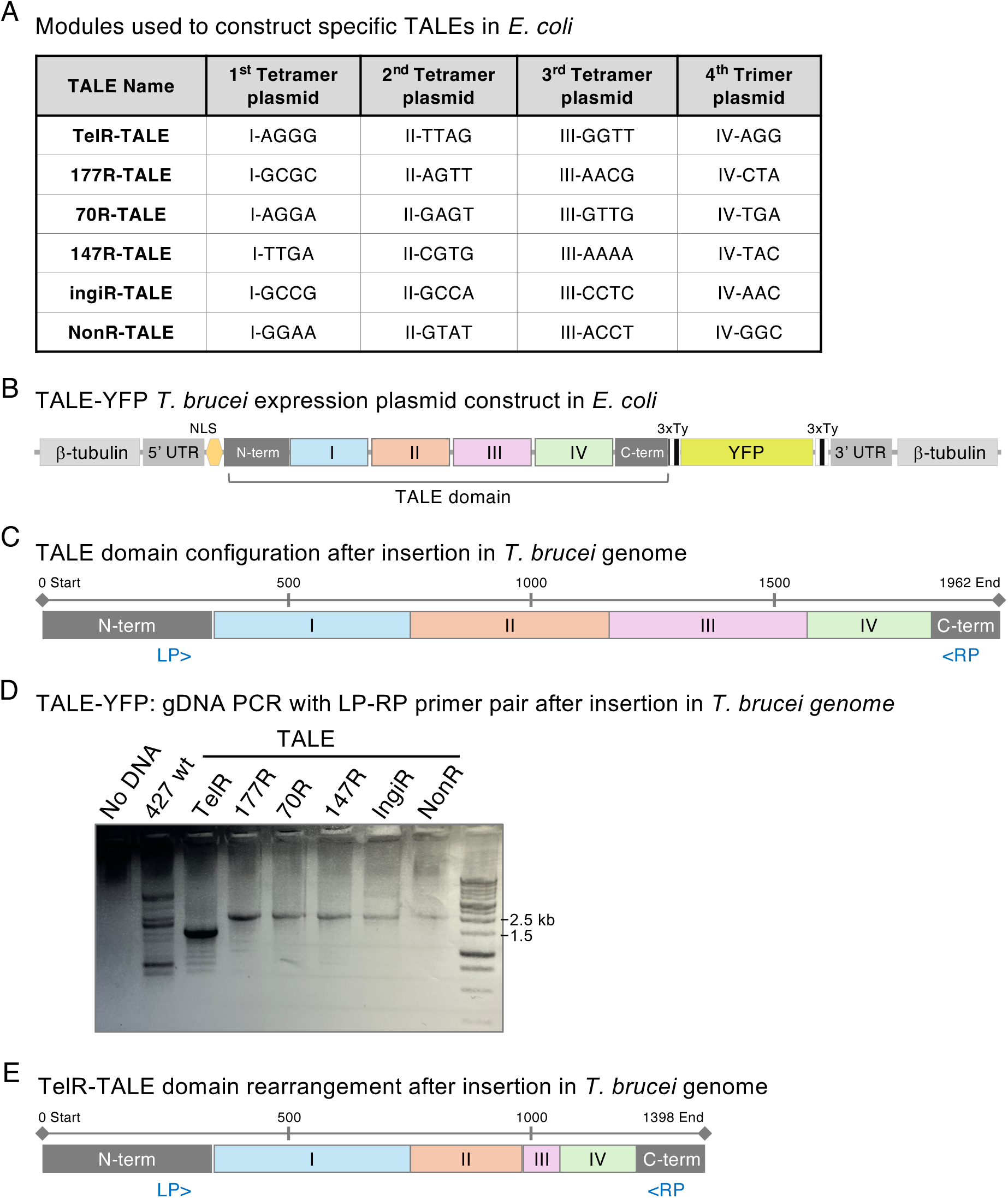
TALE-YFP construction and sequencing reveals rearranged TALE domain in TelR-TALE-YFP following intergration in the *T. brucei* genome. **A.** Tetramer and trimer modules used to build plasmids designed to express each of six TALE-YFP when integrated in the *T. brucei* genome. **B.** Each of the complete TALE plasmid contructs are expected to contain four modules comprising a complete TALE domain of the same size in all. **C.** Sequencing of five assembled TALE-YFP constructs revealed that they had the expected layout following integration at the β-tubulin locus in *T. brucei*. **D.** PCR of genomic DNA extracted from 427 control cells and 427 cells with the TelR-TALE, 177R-TALE, 70R-TALE, ingiR-TALE or NonR-TALE constructs integrated at the β-tubulin locus confirms correct size (2 kb). PCR product for TelR-TALE is shorter than expected (1.5 kb). Position of Left (LP) and Right (LP) primer pair, common to all TALE-YFP constructs, are indicated. **E.** Following integration at the β-tubulin locus sequencing revealed that the TALE DNA binding domain of TelR-TALE had rearranged explaining the shorter TelR-TALE protein made by *T. brucei* 427 TelR-TALE expressing cells (Figure S2).

**Figure S2.**
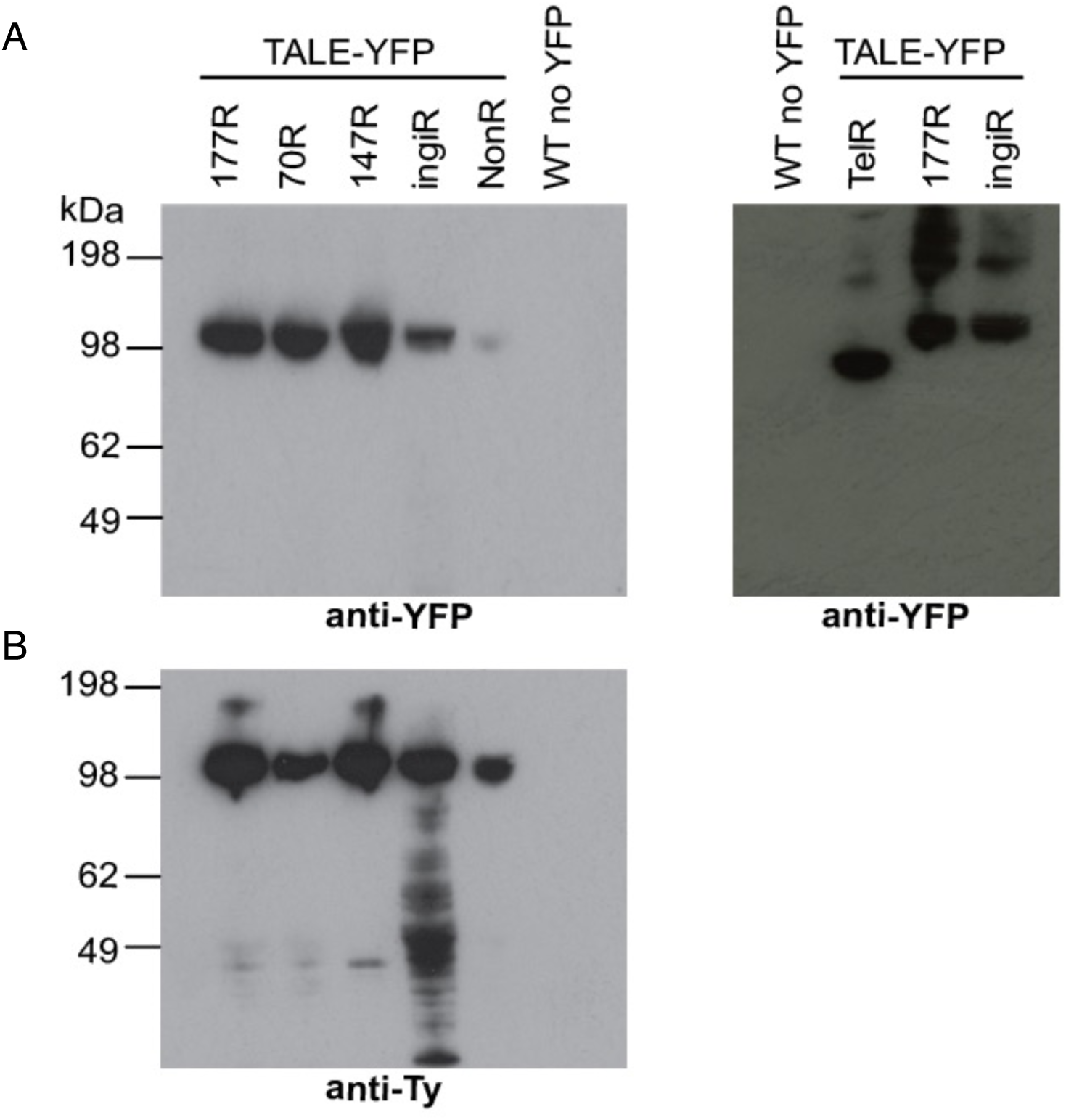
Synthetic TALE proteins are expressed in Lister 427 *T. brucei* bloodstream form cells but the TelR-TALE protein is shorter than expected. Protein extracted from 427 cells and 427 cells with constructs designed to express 177R-TALE, 70R-TALE, 147R-TALE, ingiR-TALE, NonR-TALE and TelR-TALE fused to Ty and YFP tags integrated at the β-tubulin locus was subject to western analysis using either: **A.** monoclonal mouse anti-GFP (anti-YFP) or **B.** anti-BB2 (anti-Ty). The TelR-TALE protein is smaller than the 177R-TALE, 70R-TALE, 147R-TALE, ingiR-TALE and NonR-TALE proteins.

**Figure S3.**
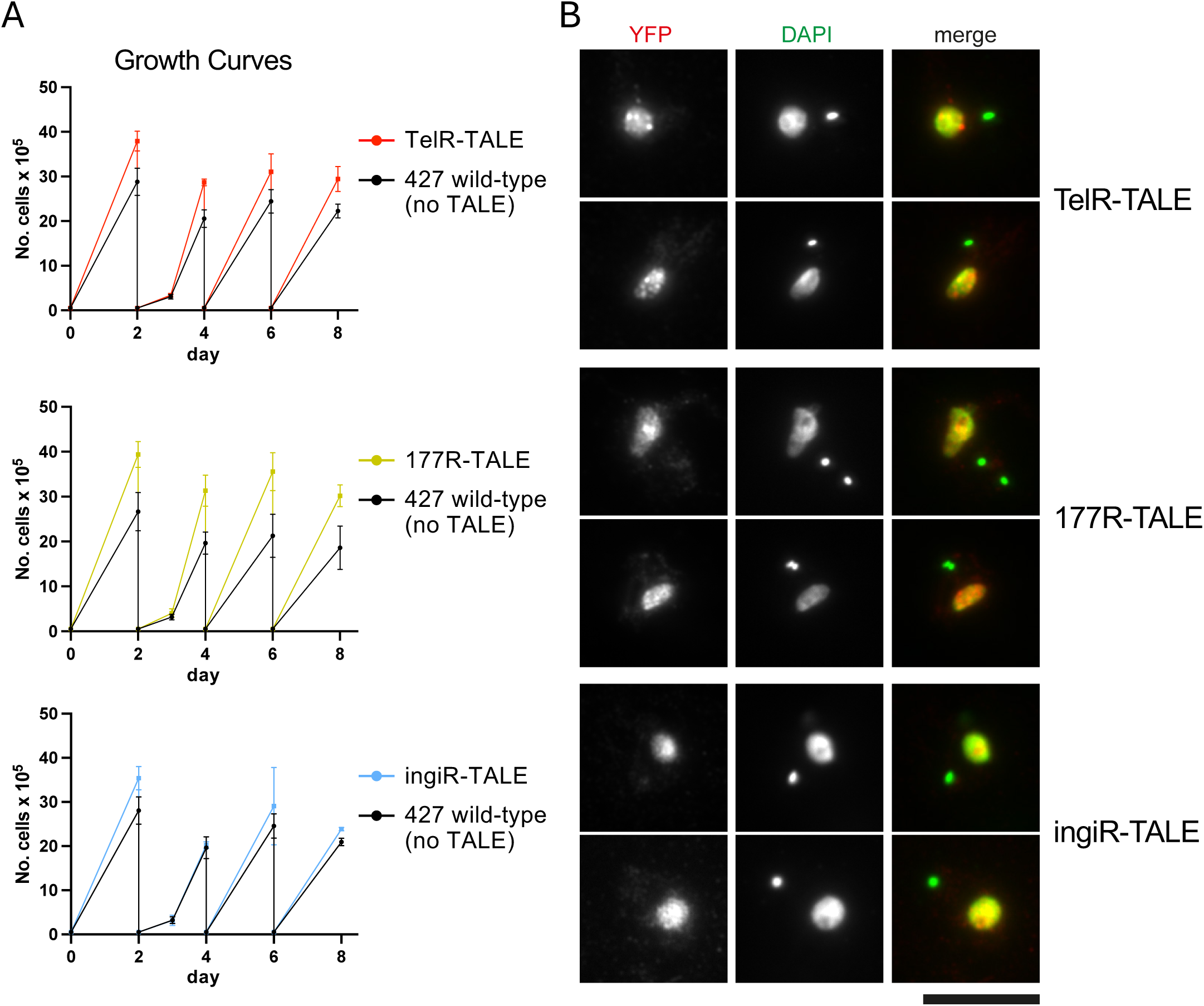
Growth assays of cells expressing TelR-TALE-GFP, 177R-TALE-GFP or ingi-TALE-GFP and their cellular localisation. **A.** Indicated cell cultures were seeded, cell number monitored and diluted every two days. **B.** DAPI and anti-GFP staining of fixed cells expressing indicated synthetic TALE-GFP fusion proteins. Bar = 10μm.

**Figure S4.**
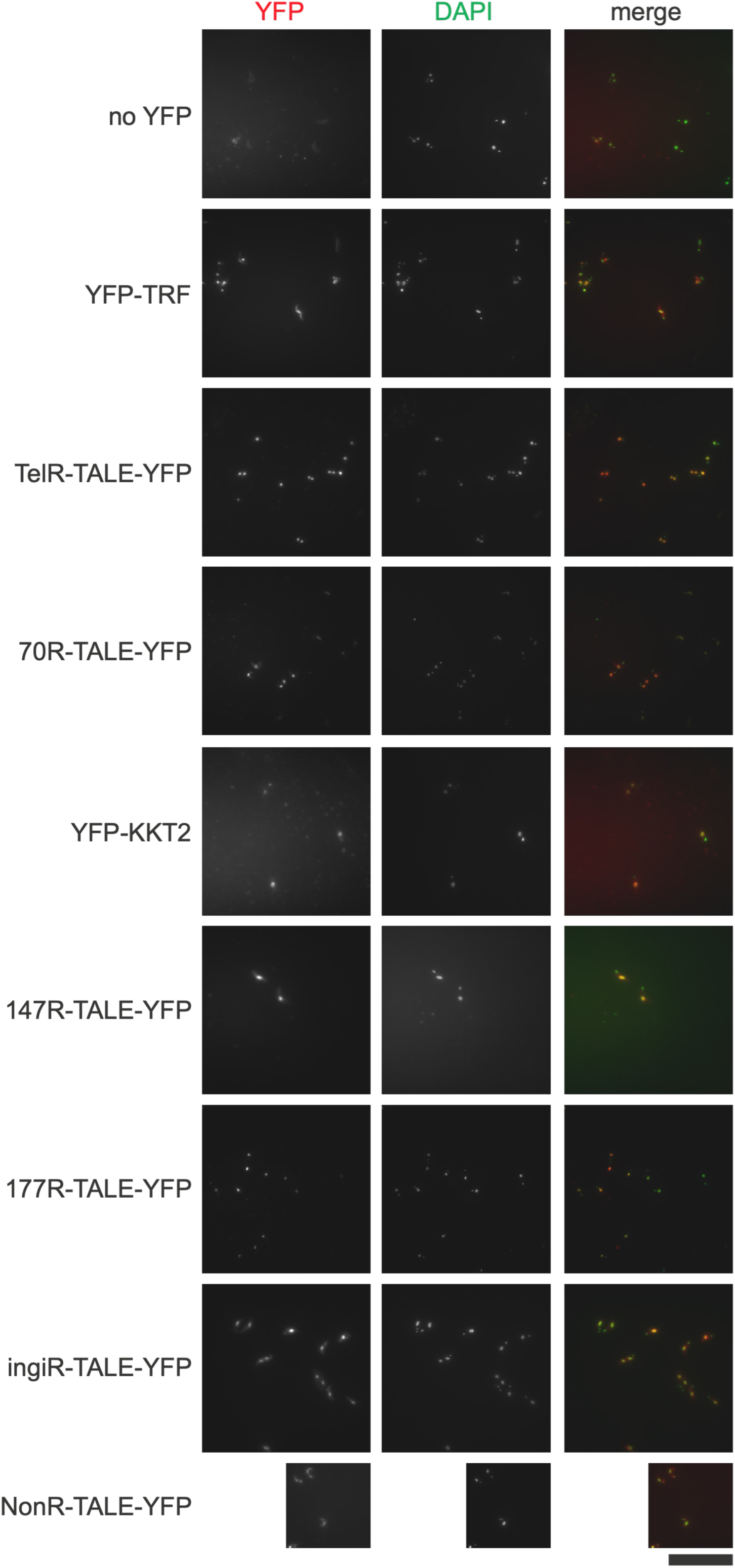
Fields of *T. brucei* cells showing the cellular localization of six expressed synthetic TALE-YFP fusion proteins compared to YFP-TRF and YFP-KKT. Bloodstream form Lister 427 *T. brucei* cells expressing the indicated TALE-YFP fusion proteins fixed and TALE-YFP protein localization detected with anti-GFP primary antibody and Alexa Fluor 568-labeled secondary antibody (red). Nuclear and kinetoplastid (mitochondrial) DNA were stained with DAPI (green). Controls cell expressing telomeric YFP-TRF, centromeric YFP-KKT2 kinetochore protein or wild type Lister 427 cells expressing no YFP, are also shown. Scale bar, 10 μm. NonR-TALE-TFP field was captured at a quarter the size of others.

**Figure S5.**
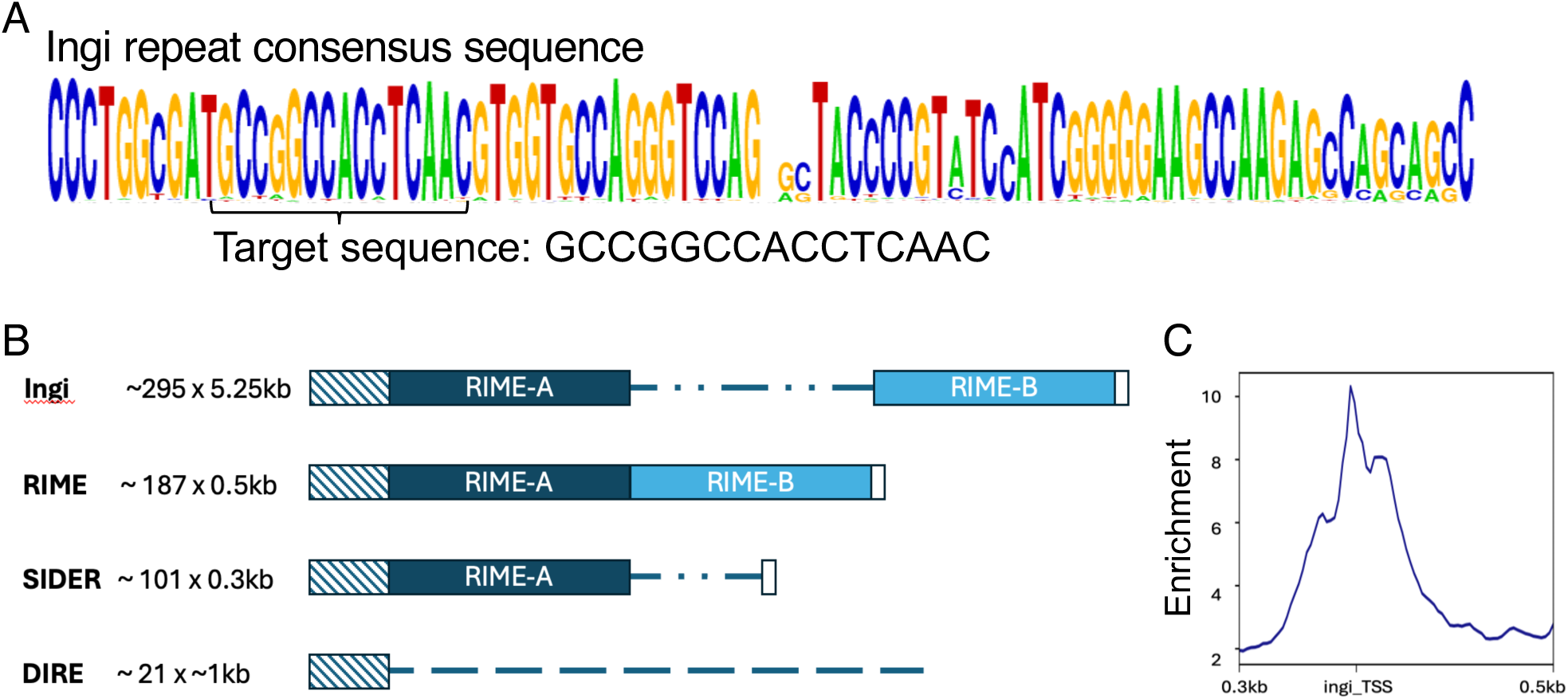
IngiR-TALE is enriched at matching binding sites located in retrotransposons. **A.** ‘Ingi’ repeat consensus sequence conserved in Ingi, RIME, SIDER and DIRE elements. The sequence that ingiR-TALE was designed to bind is indicated. **B.** Cross-hatched rectangle indicates the conserved region in Ingi, RIME, SIDER and DIRE retrotransposons, which are predicted to provide approximately 295, 187, 101 and 21 binding sites for the ingiR-TALE, respectively. **C.** Analysis of ChIP-seq data for cells expressing ingi-TALE shows enrichment of DNA residing at or near predicted ingiR-TALE binding sites.

**Figure S6.**
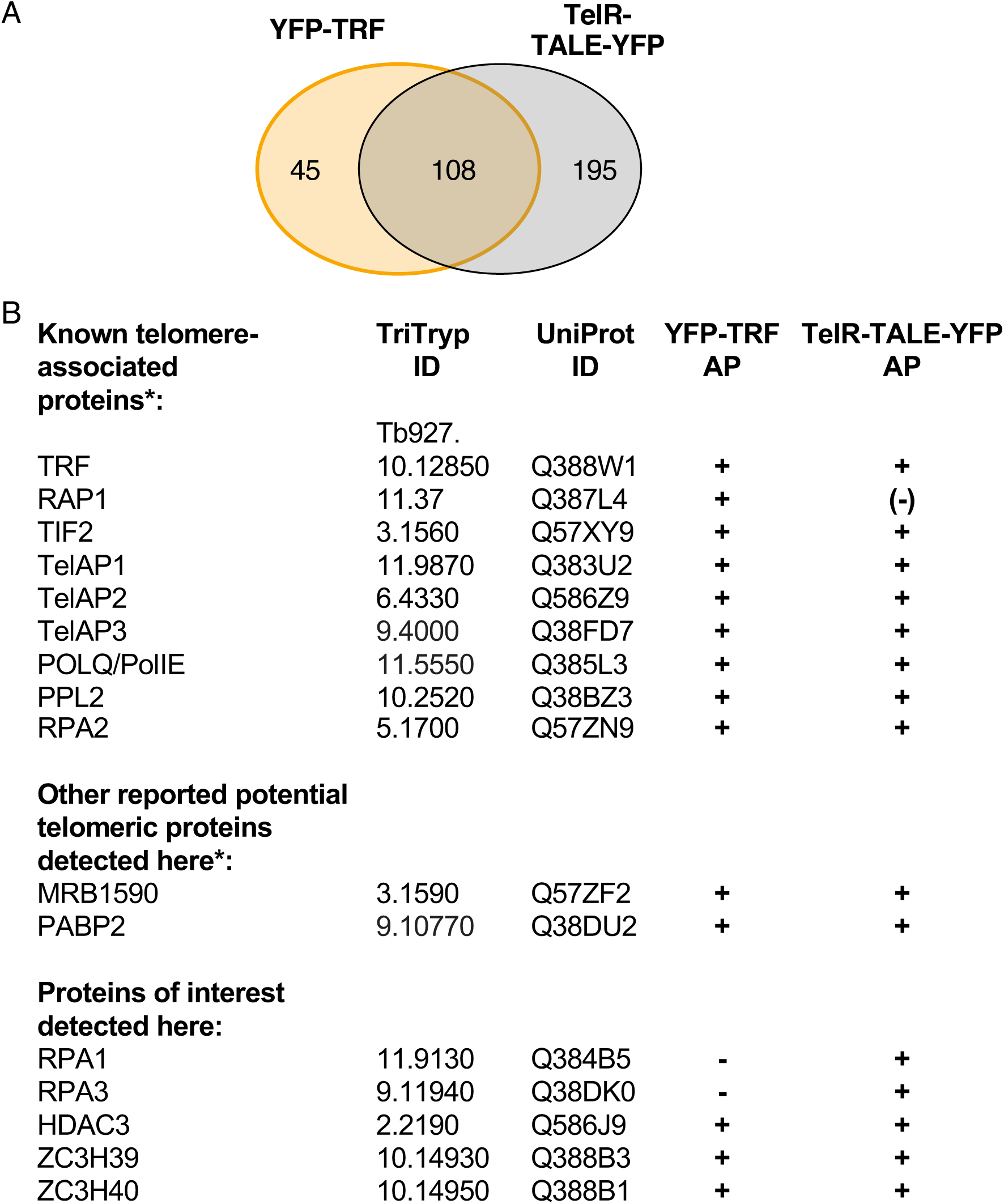
Overlap of proteins enriched in affinity purifications of both synthetic protein telomere binding protein YFP-TRF and TelR-TALE-YFP. **A.** The <u>T</u>elomere <u>R</u>epeat binding <u>F</u>actor TRF binds (TTAGGG)_n_ repeats at the ends of *T. brucei* chromosomes. A Venn diagram is shown comparing number of proteins enriched in YFP-TRF versus TelR-TALE-YFP affinity purifications. **B.** List of known telomere associated porteins and other proteins detected in YFP-TRF and/or TelR-TALE-YFP affinity purifications. + detected, - not detected, (-) weakly detected. Lists of all proteins detected in YFP-TRF and TelR-TALE-YFP affinity purifications are available in Tables S1 and S2, respectively. *See Reiss et al, 2018; Leal et al, 2020; Weisert et al, 2024.

**Figure S7.**
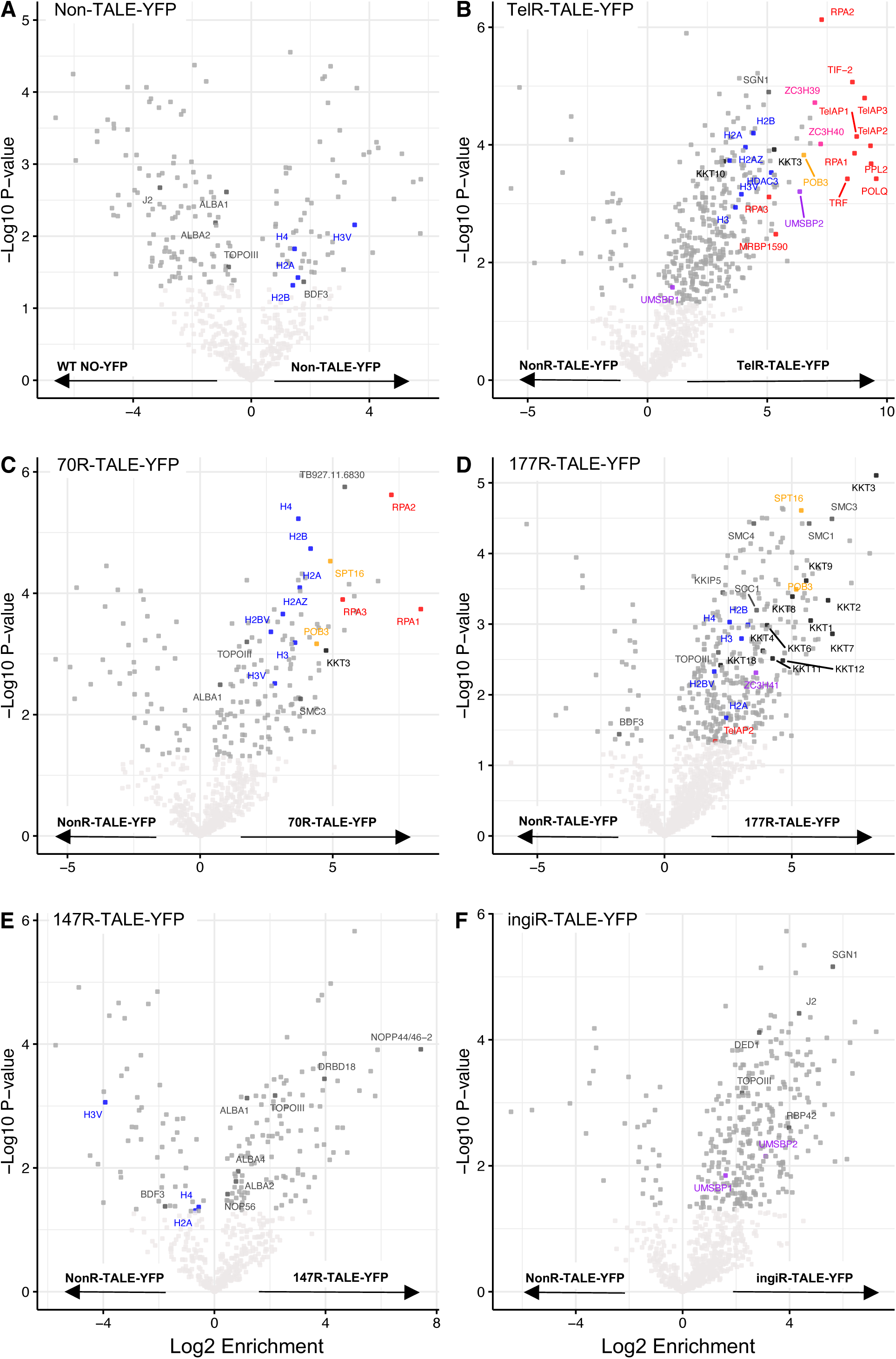
A control TALE that binds no specific *T. brucei* sequence validates proteins enriched in TelR-TALE, 70R-TALE and 177R-TALE affinity purifications. A control NonR-TALE was designed to bind the sequence GGAAGTATACCTGGC that is not present in the *T. brucei* 427 genome. Affinity selection was performed on cells expressing the synthetic NonR-TALE-YFP, TelR-TALE, 70R-TALE-YFP, 177R-TALE-YFP, 147R-TALE-YFP or ingiR-TALE proteins and control cells expressing no-YFP tagged protein. Proteins enriched with the five repeat sequence targetted TALE-YFP proteins were identified and quantified by LC-MS/MS analysis relative to the NonR-TALE-YFP control rather than the No-YFP tag control. The data for each plot is derived from three biological replicates. Cutoffs used for significance: log_2_(tagged/untagged) *P*< 0.05 (Student’s *t*-test). Enrichment scores for proteins identified in each affinity selection are presented in Supplementary Tables (Excel files).

**Figure S8.**
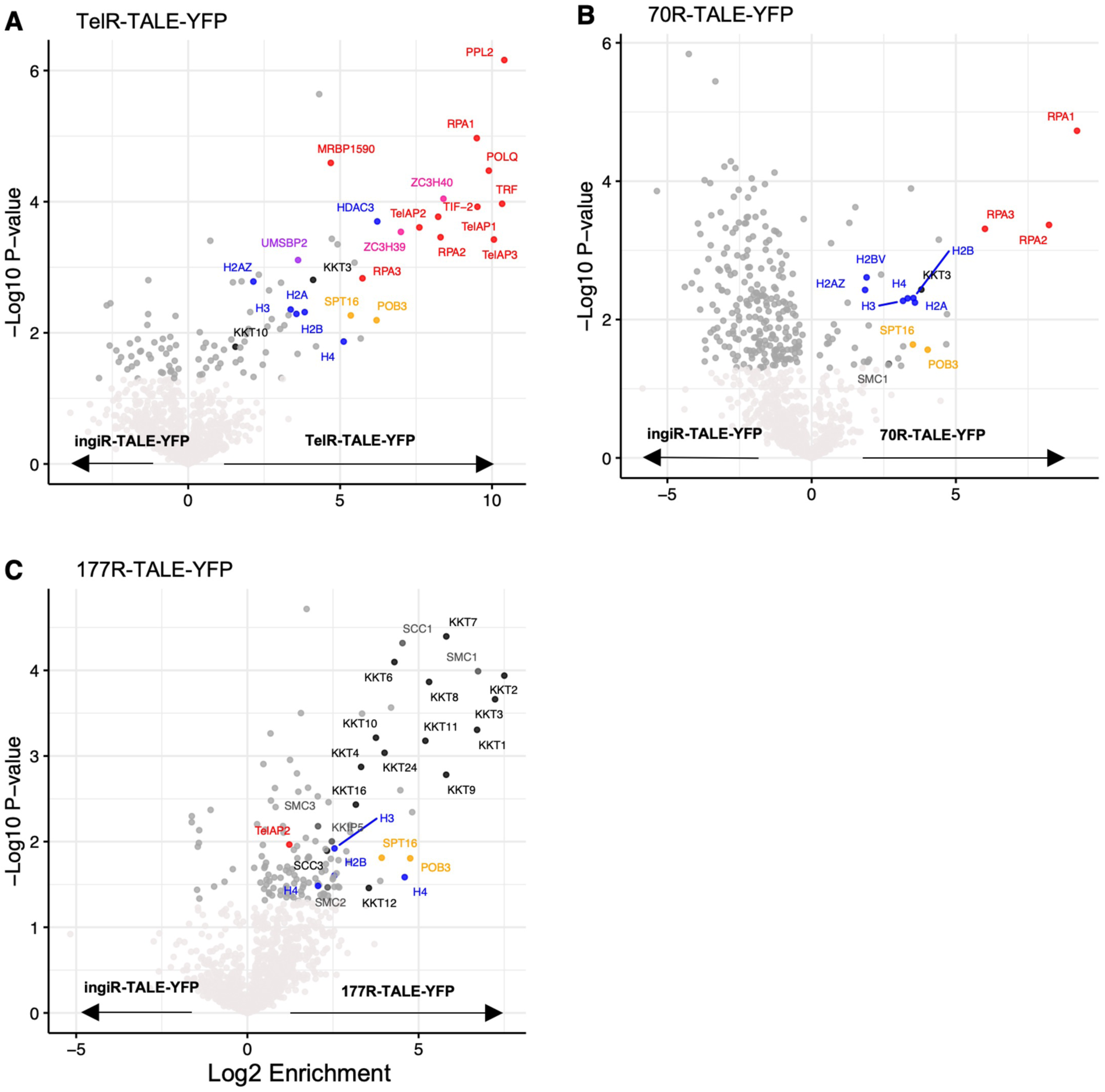
Affinity selection of TelR-TALE-YFP, 70R-TALE-YFP 177R-TALE-YFP relative to ingiR-TALE-YFP validates specificity. Affinity selection was performed on cells expressing synthetic TelR-TALE-YFP (**A**), 70R-TALE-YFP (**B**), 177R-TALE-YFP (**C**). Enriched proteins were identified and quantified by LC-MS/MS analysis relative to affinity selected ingiR-TALE-YFP as a negative control. The data for each plot is derived from three biological replicates. Cutoffs used for significance: *P*< 0.05 (Student’s *t*-test). Enrichment scores for proteins identified in each affinity selection are presented in Supplementary Tables (Excel files).

**Figure S9.**
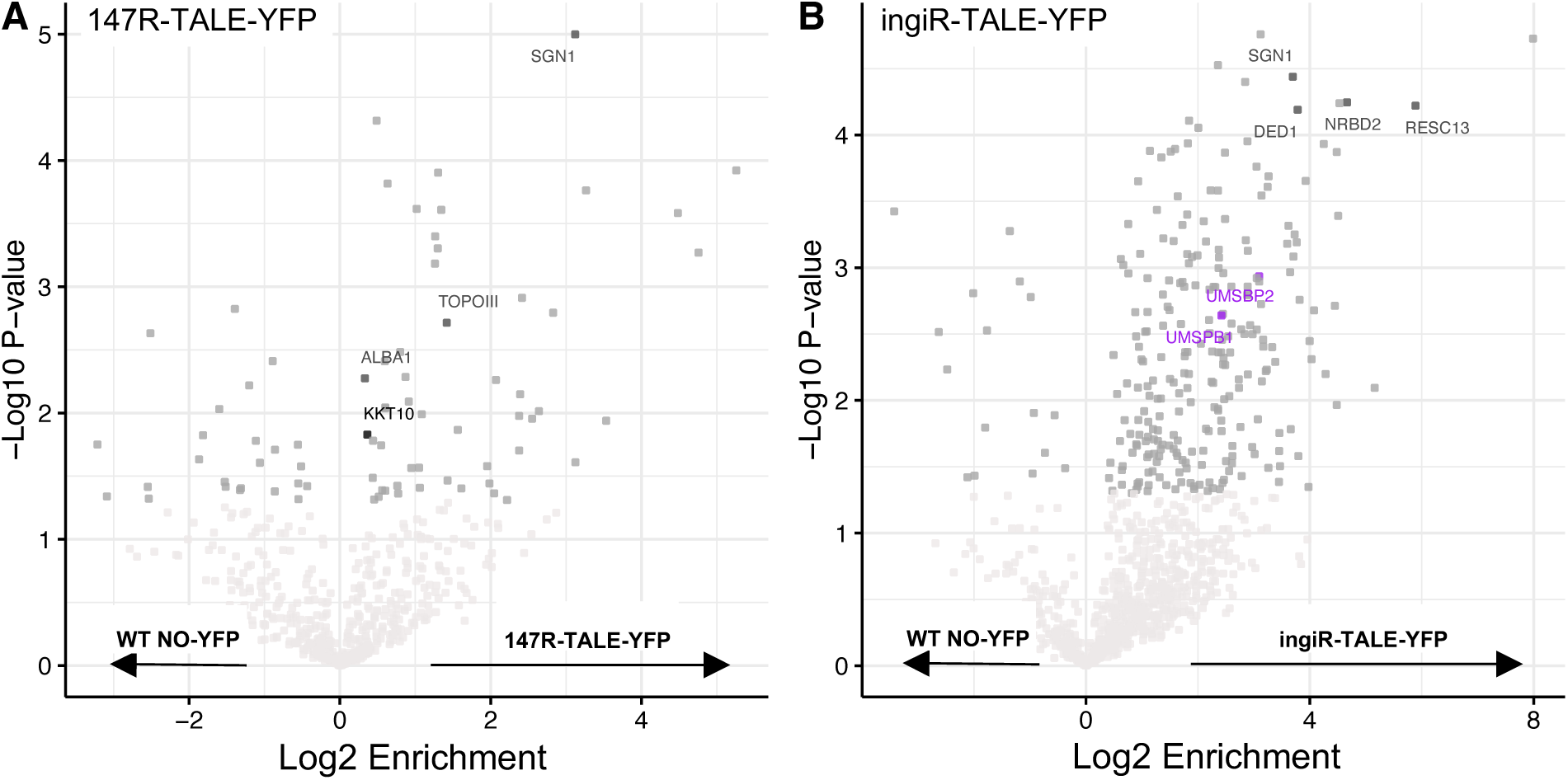
No proteins of interest are detected following affinity selection of 147R-TALE or ingiR-TALE. Affinity selection was performed on cells expressing synthetic (**A**) 147R-TALE-YFP or (**B**) ingiR-TALE-YFP proteins and control cells expressing no-YFP tagged protein. Enriched proteins were identified and quantified by LC-MS/MS analysis relative to the No-YFP tag control. The data for each plot is derived from three biological replicates. Cutoffs used for significance: log_2_(tagged/untagged) *P*< 0.05 (Student’s *t*-test). Enrichment scores for proteins identified in each affinity selection are presented in Supplementary Tables (Excel files).

**Figure S10.**
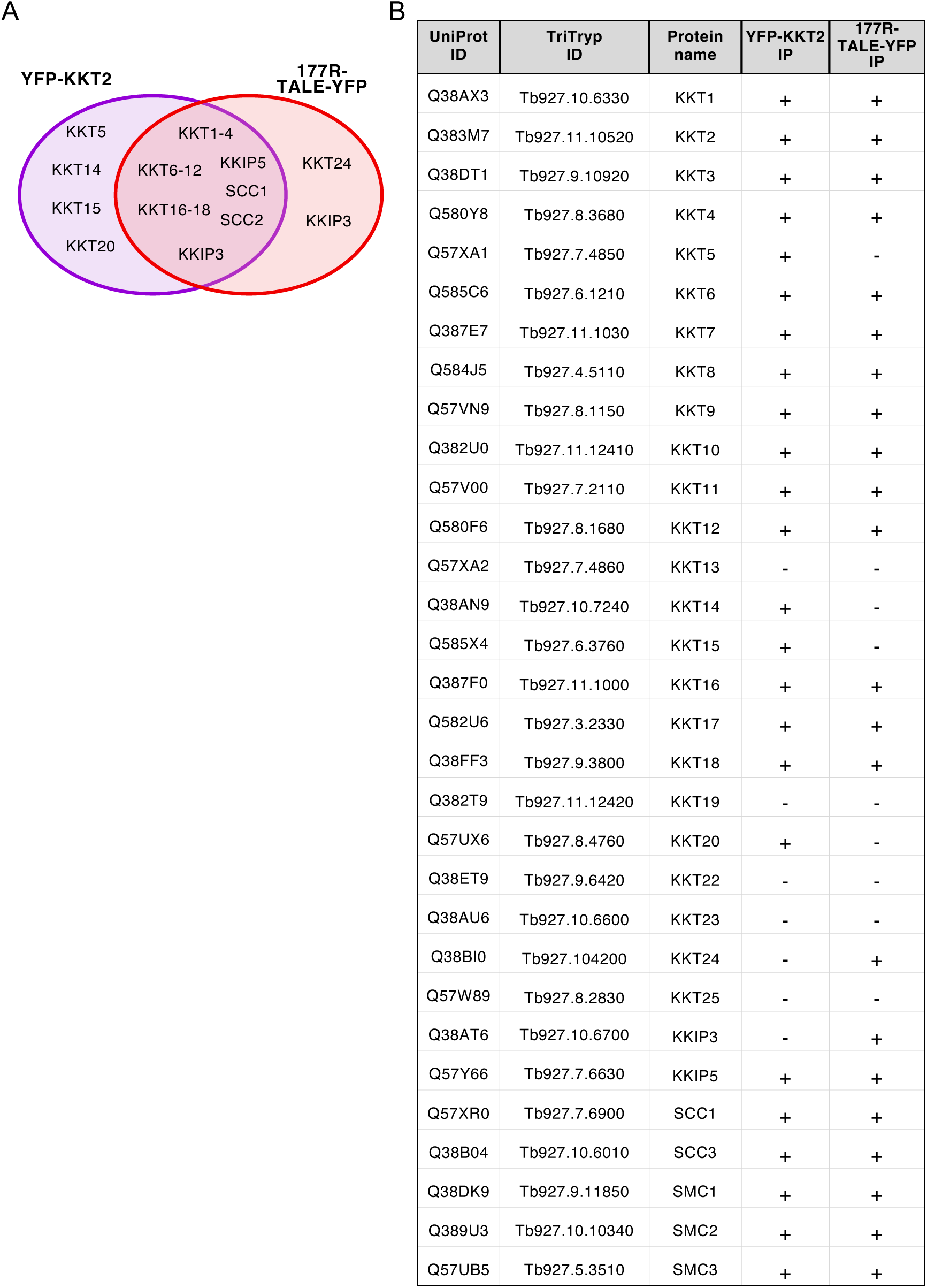
Overlap of proteins enriched in affinity purifications of both kinetochore protein YFP-KKT2 and synthetic protein 177R-TALE. <u>K</u>inetoplastid <u>K</u>ine<u>T</u>ochore (KKT) proteins are known to be enriched at all centromeres on *T. brucei* main chromosomes (Akiyoshi and Gull 2014). 177bp repeats are confined to intermediate-sized or mini-chromosomes. **A.** Venn diagram comparing proteins enriched in YFP-KKT versus 177R-TALE versus affinity purifications. A high proportion of KKT proteins, in addition to cohesin (SCC1, SCC3, SMC1 and SMC3) and condensin (SMC2) subunits, are enriched on 177 bp repeats. **B.** List of kinetochore, cohesin and condensin proteins detected in YFP-KKT and/or 177R-TALE affinity purifications. Lists of all proteins detected in 177R-TALE-YFP and YFP-KKT2 affinity purifications are available in Tables S12 and S14, respectively.

